# Sustained photoprotection involves enhanced fluorescence intermittency in a subpopulation of LHCII

**DOI:** 10.64898/2026.08.24.746725

**Authors:** Aurélie Crepin, Madeline P. Hoffmann, Cristian Ilioaia, Edel Cunill-Semanat, Andrew Pascal, Bruno Robert, Elisabet Romero, Gabriela S. Schlau-Cohen, Alizée Malnoë

**Affiliations:** Umeå Plant Science Centre (UPSC), Department of Plant Physiology, Umeå University, Umeå, Sweden; Aix-Marseille Univ, CEA, CNRS, Institute of Bioscience and Biotechnology of Aix-Marseille (BIAM), LGBP Team, Marseille, France; Department of Chemistry, Massachusetts Institute of Technology, Cambridge, MA, USA; SB2SM, CEA-Saclay, France; Institute of Chemical Research of Catalonia (ICIQ), The Barcelona Institute of Science and Technology, Tarragona, Spain; Department of Biology, Indiana University Bloomington, IN, USA

**Author notes:** These authors contributed equally to this work (Aurélie Crepin, Madeline P. Hoffmann).

**Keywords:** Photosynthesis, Photoprotection, Non-Photochemical Quenching, Light-Harvesting Complex II, Single-Molecule, Fluorescence Intermittency

## Abstract

Photoprotection against excess energy is essential for the survival of photosynthetic organisms under adverse conditions. In plants, excess energy can be dissipated as heat through non-photochemical quenching (NPQ) of chlorophyll fluorescence, involving the trimeric light-harvesting complex II (LHCII), the major antenna of photosystem II. How NPQ affects antenna proteins remains debated, especially as most studies focus on short-lived components artificially induced in vitro. Here, we characterize the effects of qH, a long-lived NPQ component, on the fluorescence properties of natively quenched LHCII. Single-molecule fluorescence measurements, combined with biochemical and biophysical ensemble approaches, reveal a larger and more quenched subpopulation of LHCII trimers exhibiting fluorescence intermittency in samples with qH compared to those without. This behavior is linked to a small conformational change that stabilizes a quenched state, enhancing photoprotection at the antenna level. These findings provide new insights into sustained NPQ and its role in regulating energy dissipation under natural light conditions.

**Teaser:** Enhancement of a quenched, blinking state in a subpopulation of LHCII trimers characterizes sustained photoprotection in plants.

## INTRODUCTION

An essential feature of photosynthetic organisms is protection against damage from excess absorbed energy under conditions where the electron transport chain becomes saturated. This protection is particularly important under abiotic or biotic stress conditions that decrease metabolic efficiency (*1*). The current context of rising temperatures and increasing frequency of climate events, such as heat and drought waves, means there is an urgent need to understand how these organisms protect themselves from excess light, how to fine-tune such protection for increased climate resilience, and how to incorporate natural strategies into artificial systems (*2*, *3*).

Some photoprotective pathways are characterized by the safe dissipation of excess energy as heat. These pathways are detected as a decrease in chlorophyll (Chl) fluorescence when photochemistry is saturated by a strong light pulse and are therefore called non-photochemical quenching (NPQ) of Chl fluorescence. NPQ comprises different components defined by their effectors and by the kinetics of activation and relaxation (for a review, see, e.g. (4, 5)). Several of these components have a common target: the light-harvesting antennae proteins, the pigment-protein complexes involved in energy transfer to the photosystems core. The most abundant type of antenna protein in plants is the light-harvesting complex II (LHCII), the major antenna of photosystem II (PSII). LHCII takes the form of a membrane-embedded heterotrimer of various combinations of Lhcb1, Lhcb2 and Lhcb3 subunits and isoforms (*6*). Each LHCII trimer contains a total of 42 chlorophyll molecules (24 Chl *a* and 18 Chl *b*) as well as 12 xanthophylls, a subclass of carotenoids (Car) (for each monomer on average: one violaxanthin or zeaxanthin (Vio, Zea), one neoxanthin (Neo) and two luteins (Lut1, Lut2)) scaffolded by three transmembrane helices (A, B and C) within each monomer (*7*, *8*). Like many known antenna proteins, LHCII trimers are capable of both light harvesting and energy dissipation (*9*), most likely via conformational changes in the protein scaffold (*10–14*).

The leading model for conformational change-induced quenching in LHCII includes the repositioning of the lumenal loops and a decrease in the crossing angle of the transmembrane helices A and B (*15*, *16*). The latter would bring closer Lut1 and the terminal emitter Chl pair, *a*611 and *a*612 (*11*, *17–19*), as was observed in lipidic environment where LHCII is more constrained (*10*, *16*). Distortions in Lut1 and Neo were revealed by resonance Raman vibrational spectroscopy in LHCII particles quenched by detergent removal in polyacrylamide gels, which limits protein aggregation (*20*). Beyond single-particle processes, antenna reorganization, and in particular LHCII aggregation, have also been suggested to drive quenching (*10*, *21–27*). However, the site(s) and mechanism(s) of quenching in LHCII remain a subject of discussion and investigation as several mechanisms may co-occur or occur in different conditions (see (*28–30*) for recent reviews).

One of the reasons for the ongoing debate around quenching in LHCII is that most of these studies characterize LHCII quenching in the context of energy-dependent quenching (qE), which relies in plants on the acidification of the lumen pH, the activation of antenna-like PsbS protein and/or the conversion of Vio to Zea (*23*, *31–35*). qE is activated and relaxed in seconds to minutes, preventing isolation of natively quenched systems. Yet, as thylakoid membranes are highly packed with photosystems and antennae of similar pigment composition (*36*), the identification of quenching site on specific antennae has to be performed in vitro. The study of LHCII in the context of qE thus relies on induction of quenching post-purification, by placing LHCII trimers in conditions such as PsbS-containing liposomes, increased lateral pressure from membrane mimetics, acidic buffers and/or aggregation by detergent removal, see e.g. (*10*, *16*, *24*, *37–41*). Several studies have reported similarities between these in vitro samples and in vivo measurements (e.g. (42–45)), yet it can still be difficult to define physiologically relevant in vitro conditions (46–48). Differences in these conditions may contribute to the diversity in models and thus the controversy over the quenching sites and mechanisms.

Our recent work revealed that LHCII trimers also serve as a site for the sustained NPQ component, qH, in *Arabidopsis thaliana* (*49*). This pathway is induced by the chloroplast lumen-localized protein LIPOCALIN IN THE PLASTID (LCNP), without which NPQ is decreased and plant photobleaching increased under strong stress such as cold and high light combination (*50*). qH is also regulated by the proteins SUPPRESSOR OF QUENCHING 1 (SOQ1) and RELAXATION OF QH 1 (ROQH1), which respectively modulate the activation and relaxation of qH (*51–53*) (Figure 1A). The mutant of the two negative regulators, *soq1 roqh1*, exhibits stunted growth due to constitutive energy dissipation (i.e. occurring under standard growth light conditions), which limits light availability for photochemistry; this phenotype is reversed by knocking out LCNP (*52*). Interestingly, qH is independent of PsbS, lumenal pH changes, or Vio, Zea, or Lut (49–52), distinguishing its activation from qE. The working model for qH modulation is redox-based rather than through protonation: under non-stress conditions, qH is prevented by SOQ1 through its methionine sulfoxide reductase activity on LCNP; under stress conditions, such as cold and high light, LCNP methionines become oxidized, thereby activating LCNP for the formation of qH at the thylakoid membrane in the major, as well as minor, antenna of PSII (*53*, *54*). LCNP is a soluble protein, whose expression is increased during abiotic stress such as drought or high light (*55*). Based on domain conservation, LCNP is an eight-stranded antiparallel beta sheet that forms a barrel with high affinity for small hydrophobic molecules. Under excess light conditions, LCNP would modify a molecule in the vicinity of or within the antenna proteins, thereby triggering a conformational change that converts antenna proteins from a light harvesting state to a dissipative state (*4*).

**Fig. 1.**
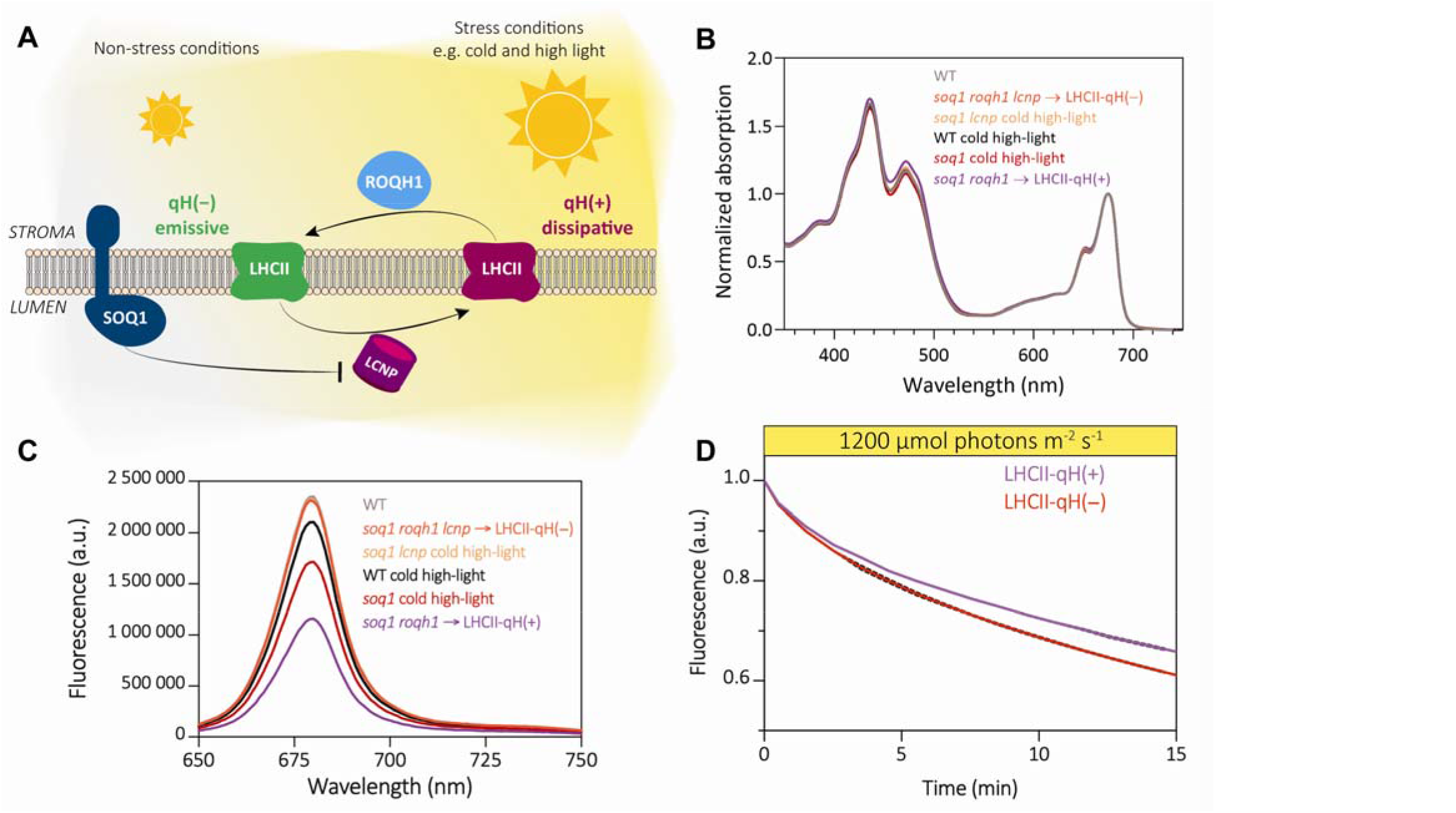
*soq1 roqh1* LHCII are natively quenched, and this quenching is photoprotective. **(A)** Simplified scheme of the qH mechanism. LCNP, a lumenal protein, is essential for qH activation in stress conditions and energy dissipation from antenna proteins. LCNP is inhibited by SOQ1. ROQH1, a stromal protein, allows qH relaxation. (**B)** Normalized room temperature absorption spectra of wild type and mutant LHCII trimers. Spectra are normalized to the peak at 675 nm. (**C)** Room temperature fluorescence emission spectra of wild type and mutant LHCII trimers, normalized on the maximum sample absorption in the Qy peak, upon excitation at 625 nm. **D)** Normalized steady-state fluorescence of purified LHCII trimers over a 15 min incubation under high light to induce photobleaching. Data represent mean ± SD of three biological repeats (from different thylakoid preparations) and normalized on initial fluorescence of the sample.

Due to the recent discovery of qH, it remains to be deciphered if this quenching component relies on a different regulation of a similar energy dissipation pathway to qE, or on different pigment-protein configurations entirely. Unlike qE, the slower kinetics (minutes to hours) and pH-independent properties of qH provide a unique opportunity to isolate and perform in-depth characterization of natively and stably quenched LHCII without artificial induction. Previously we determined that qH is also insensitive to the subunit composition of LHCII and that pigment and lipid composition in quenched vs unquenched trimers were similar (although a small modification in a few trimers may have escaped detection) (*49*). The molecular origin for quenching is thus an outstanding question that we aim to answer.

Here, we evaluate the structural and functional properties of natively quenched LHCII trimers purified from the *soq1 roqh1* double mutant with active qH under non-stress conditions. Using chlorophyll fluorescence measurements at both the ensemble and the single-molecule levels, we demonstrate that qH correlates with an enhanced propensity for fluorescence intermittency (“blinking”) that gives rise to quenching in a subpopulation of LHCII particles, independently of trimer-trimer interaction. Using circular dichroism spectroscopy and resonance Raman spectroscopy, we propose that a small conformational change, possibly in the helix C region of the Lhcb proteins and its neighboring chlorophyll *b* cluster, could explain this behavior. These results not only identify key features of direct photoprotection in antenna proteins, but also open new avenues for modelling, manipulating and mimicking photoprotection *in vivo*.

## RESULTS

### Native LHCII from the *soq1 roqh1* mutant display large fluorescence quenching

LHCII trimers were purified using fast protein liquid chromatography (FPLC) from wild type (WT) Arabidopsis plants and four mutants, with higher or no capacity for qH. Plants were grown under non-stress conditions before either being harvested directly (hereafter referred to by genotype name alone) or subjected to a cold (4□) and high light (1,600 µmol photons m^−2^ s^−1^) stress for 6 h (hereafter ‘cold high-light’). These mutants lack the qH negative regulators SOQ1 and/or ROQH1 (*soq1, soq1 roqh1*) or additionally the qH activator LCNP (*soq1 lcnp, soq1 roqh1 lcnp*); by convention, lower case italic denotes a mutation in that gene. Ensemble absorption and fluorescence spectra from WT and the four mutant LHCII samples show similar and expected peak structure with maximum absorption at 675 nm in the Qy peak and fluorescence emission at 679 nm (Figure 1B). Note that the slight difference in absorption spectra was inconsistent across repeats and could not be correlated to quenching.

Having confirmed successful purification and similar absorption spectra, we compared fluorescence intensities across samples. The ensemble fluorescence intensity is quenched in LHCII from WT cold high-light, *soq1* cold high-light, and *soq1 roqh1*. The isolated LHCII from *soq1 roqh1* display the highest level of fluorescence quenching (49 ± 2 % decrease compared to WT) of all the samples (Figure 1C), even though the plants were not stressed, in line with the constitutive quenching displayed by the plants under non-stress conditions (49, 52). The isolated LHCII from the corresponding mutant controls (*soq1 lcnp* cold high-light, *soq1 roqh1 lcnp*) show fluorescence intensity at a similar level to non-stress WT. The mutant controls lack the factor responsible for qH induction and so demonstrate that the observed quenching of chlorophyll fluorescence is LCNP-dependent qH. As *soq1 roqh1* LHCII displayed the highest quenching, we used this LHCII sample (hereafter LHCII-qH(+)) and the corresponding control *soq1 roqh1 lcnp* (hereafter LHCII-qH(−)) for the rest of the study.

To test that the observed quenching is photoprotective, we exposed LHCII-qH(+) and LHCII-qH(−) to high light at ∼1200 µmol photons m^−2^ s^−1^ for 15 min (Figure 1D). After treatment, both samples showed photobleaching, but LHCII-qH(+) were significantly less affected with a fluorescence loss of 34.8 ± 0.3 % compared to 39.6 ± 0.2 % for the LHCII-qH(−). The fluorescence quenching observed in LHCII trimers with active qH therefore reflects a photophysical pathway that provides photoprotection against excess light to the antenna proteins themselves.

### qH relies on a small conformational change at single-trimer level

Previous research on isolated qH-active LHCII trimers from cold high-light treated *soq1* showed that qH was not due to an evident change in pigment composition (*49*). We further tested this here by HPLC analysis of LHCII-qH(+) and LHCII-qH(−): although displaying ∼50% less fluorescence intensity, LHCII-qH(+) has the same pigment content as the control (Figure S2). We thus employed circular dichroism (CD) spectroscopy, a highly sensitive technique to probe the local environment and excitonic properties of the pigments (*56*, *57*). The main differences are observed in the relative amplitude of the excitonic bands located at approximately (+)447, (−)475 (Soret) and (+)670 nm (Chl Qy region) (Figure 2B). The (−)475 nm excitonic band most likely originates from Chl *b* and neoxanthin (Car S_2_ absorption range) molecules while the band at (+)670 nm originates from excitonic interactions involving Chl *a* (*11*, *58–60*). A recent study attributed changes in intensity around 470 nm to Chl *b*608-b609 and *b*606-b607 interactions (*61*) while another work indicated that this perturbation is due to interactions between pigments in different monomeric subunits (*59*). Interestingly, we found that, upon dissociation of the trimer into monomers by phospholipase A2 (PLA2) treatment, a significant recovery of emission occurs in LHCII-qH(+) up to ∼93 % of the control (Figure S3), implicating pigments at the monomer interface as a likely site of quenching.

**Fig. 2.**
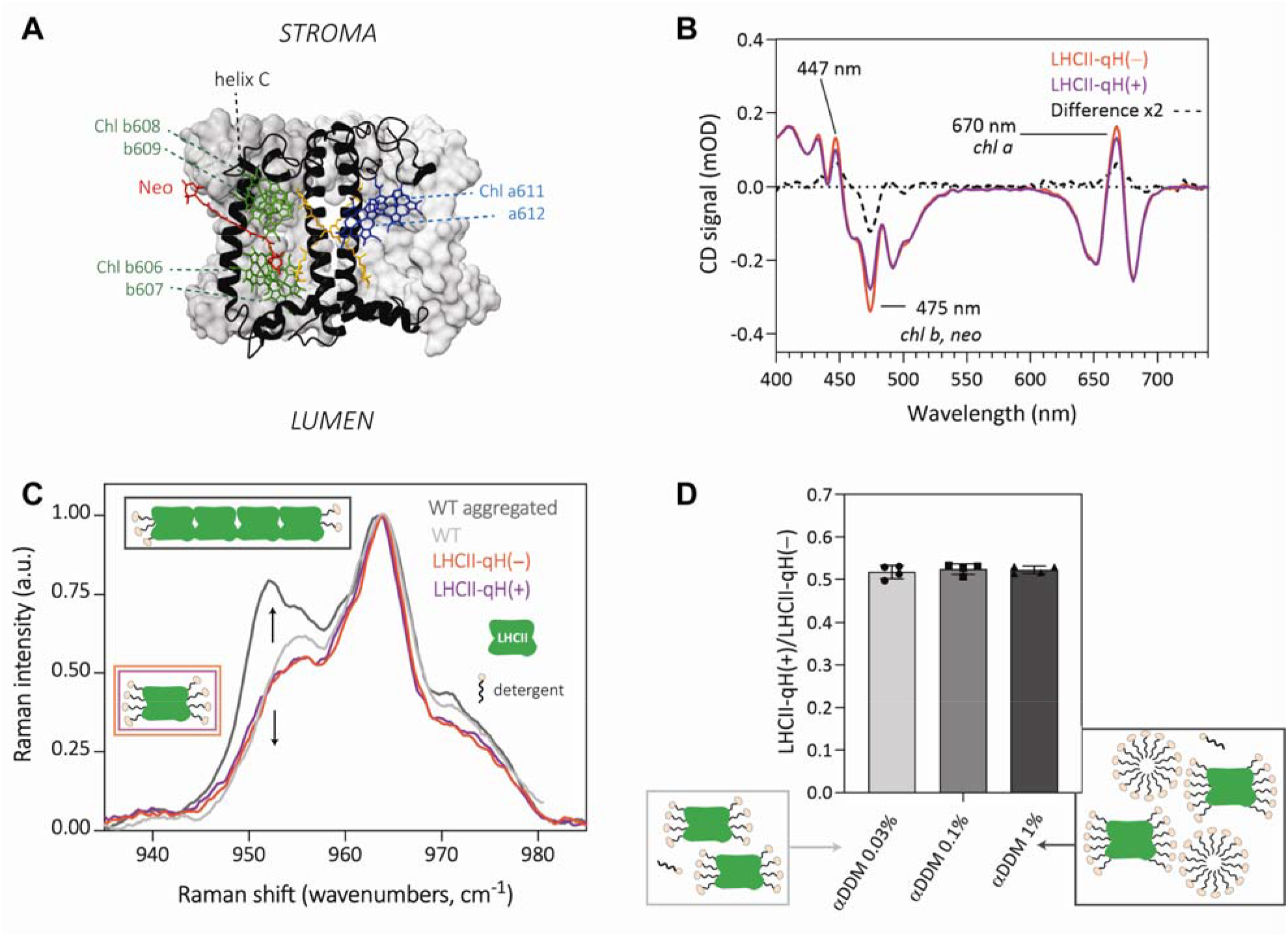
qH is likely due to a small conformational change at the single-trimer level. **(A)** Structural model of the LHCII trimer, displaying only pigments of interest: neoxanthin (Neo, red) and the two Chl *b* couples *b*608-b609 and *b*606-b607 (green), around helix C; and the terminal emitter pair a611-612 (dark blue). The structure was adapted from 8IWZ (*16*). For a structure with all pigments displayed, see Figure S1. **(B)** Circular dichroism spectra of LHCII-qH(+) (purple) and LHCII-qH(−) (orange) trimers at the same chlorophyll concentration, as well as the LHCII-qH(−) *–* LHCII-qH(+) difference multiplied by 2 (x2) (black dashed line). **(C)** ν4 region of the resonance Raman spectra of LHCII-qH(+) (purple) and LHCII-qH(−) (orange) trimers isolated in n-dodecyl β-D-maltoside (β-DDM). For comparison the spectra of untreated (light grey) or aggregated WT LHCII (dark grey) obtained by detergent removal, with a similar level of quenching than LHCII-qH(+) are displayed. Excitation was at 488 nm. **(D)** Ratio of room temperature fluorescence of LHCII-qH(+) and LHCII-qH(−) trimers incubated in an increasing a-DDM detergent concentration to break trimer-trimer interactions. Data represent mean ± SD (n=4 replicates from two different purifications on two different thylakoid preparations). Inset represents detergent conditions, with LHCII in green and detergent in beige and black.

These results thus suggest that qH induces changes in pigment environment, which could stem from protein conformational changes. Resonance Raman vibrational spectroscopy is a well-established technique for interrogating pigment-protein interactions and/or chemical properties of the pigments themselves: it was previously used to characterize LHCII aggregation, which has been hypothesized to be a mechanistic driver of qE (*21*, *22*). For an excitation of neoxanthin at 488 nm, in particular, aggregated LHCII were shown to display an increase in the intensity of the Raman vibrational mode at around ∼952 cm^−1^ relative to the one at 964 cm^−1^, entirely absent in non-aggregated trimeric LHCII (*22*) (Figure 2C). We thus used this technique to investigate changes in conformation around specific pigments and determine if a similar mechanism is at play for qH.

Resonance Raman spectra of LHCII-qH(+) and LHCII-qH(−) trimers both unambiguously differed from WT LHCII aggregated to a similar fluorescence yield as with qH (∼55%), for neoxanthin excitation at 488 nm (Figure 2C), as well as for excitation at 441 nm probing Chl *b* environment (Figure S4). Importantly, the Raman spectra from LHCII-qH(+) upon excitation at 488 nm does not show any increase in the intensity of the aggregation-specific peak at 952 cm^−1^ (Figure 2C). This result corroborates that qH is independent of trimer-trimer interaction. However, both LHCII-qH(+) and LHCII-qH(−) samples display similar spectra for the pigments probed (Figure 2C, Figure S4).

We further tested the role of trimer-trimer interactions in qH by subjecting LHCII-qH(+) and LHCII-qH(−) trimers to aggregation by detergent removal. Aggregation led to a large decrease in fluorescence yield in both samples, yet preserving and enhancing, the quenching in LHCII-qH(+) by 70% compared to control (Figure S5A,B). Additionally, LHCII-qH(+) and LHCII-qH(−) trimers displayed the same 77K steady-state fluorescence in detergent-solubilized suspension: a sharp peak at 680 nm (Figure S5C) devoid of a significant red shift, a feature typically associated with the formation of large LHCII aggregates (*43*, *62–65*). The ratio of the fluorescence at 700 nm to the value at 680 nm, F700/F680, was 0.081 ± 0.004 for LHCII-qH(−) and 0.086 ± 0.002 for and LHCII-qH(+). This result further substantiates an absence of LHCII aggregation in these samples. However, such low values for F700/F680 have also been described for low order clusters of LHCII trimers in vitro (*46*). To verify that qH does not depend on protein-protein interaction, we incubated the samples in increasing detergent concentrations to disrupt possible LHCII aggregates (*66*) and to promote homogeneous suspensions of single trimers in micelles, thereby limiting the formation of low-order clusters (*46*). Detergent concentrations from 0.03% (used during sample isolation) to 0.1% and up to 1% n-dodecyl α-D-maltoside (α-DDM) did not affect qH: the fluorescence ratio between LHCII-qH(+) and LHCII-qH(−) trimers was maintained throughout the experiment (Figure 2D).

Altogether, these results demonstrate that qH occurs at single LHCII trimer level as opposed to being induced by trimer-trimer interactions. It is possible that qH could be triggered by a small conformational change localized within the trimer. We can further hypothesize that the difficulty to detect clear conformational changes signature in the protein might be due to subpopulations of different quenching states present in the sample and lost in the average in ensemble measurements.

### LHCII-qH(+) access an overlapping range of photophysical states as LHCII-qH(−)

To determine if LHCII-qH(+) samples contain trimers in different quenching states, we performed ensemble fluorescence lifetime measurements as multiple lifetime decay components typically arise from the presence of multiple states (Figure 3A, Figures S6-S8). All samples were best fit with a two-component exponential decay. Each sample had lifetime components of approximately 3.5 ns and 0.2 ns (Figure 3B, Figure S6, Table S1), corresponding to unquenched and quenched states of the complex, respectively. The extracted timescales broadly agree with previous reports of LHCII lifetimes (*38*, *49*). The extracted average lifetimes also generally decreased as the fluorescence intensity decreased (Table S1), consistent with enhanced quenching. For example, LHCII-qH(+) had the largest amplitude of the short 0.2 ns lifetime component (35.9 ± 6.8%) and the lowest fluorescence intensity while the WT and LHCII-qH(−) samples were dominated by the unquenched states (Figure 3B).

**Fig. 3.**
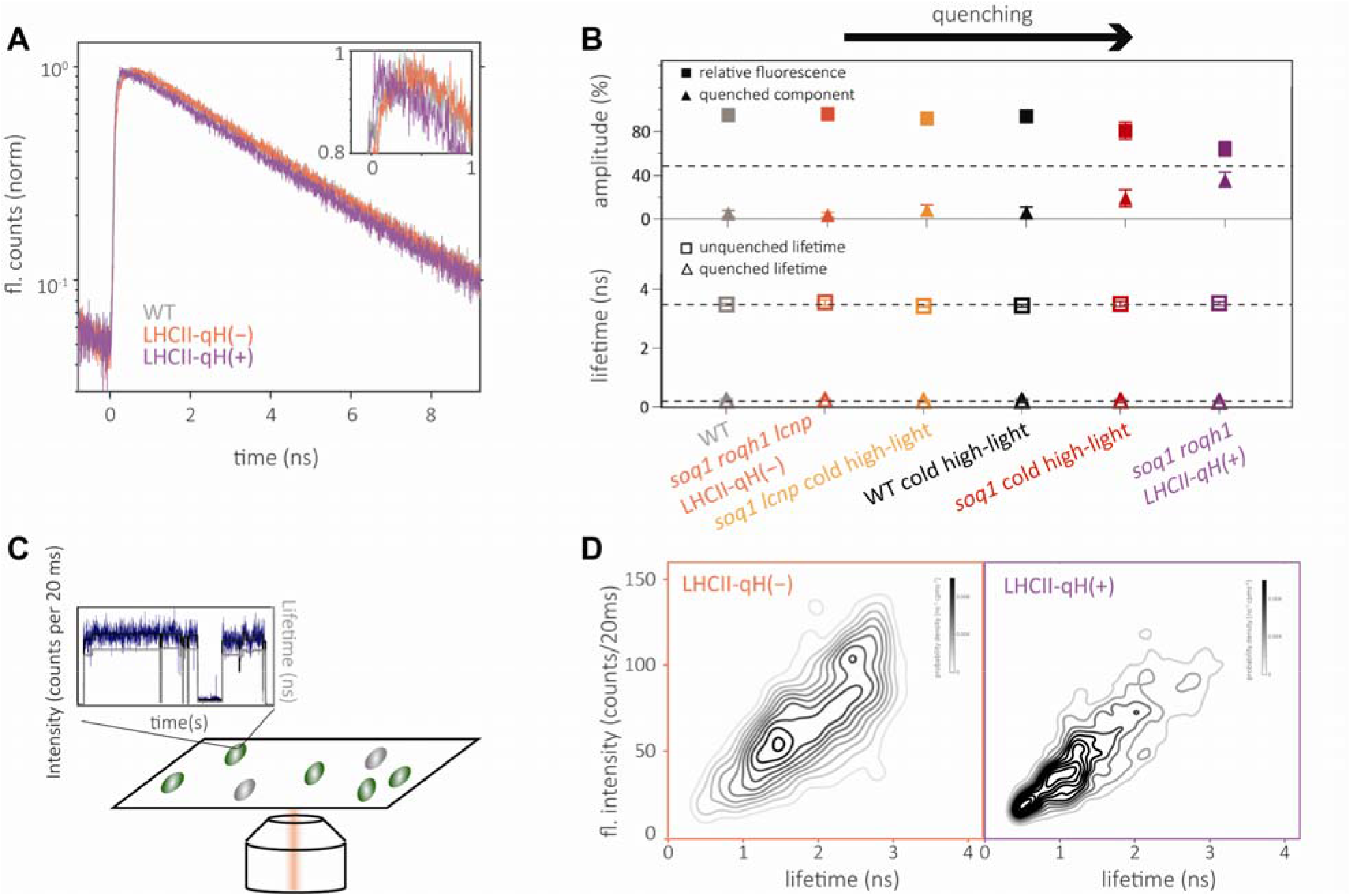
LHCII-qH(+) trimers access an overlapping range of chlorophyll excited state lifetimes with differing probabilities. **(A)** Fluorescence decay traces from time-correlated single-photon counting experiments on LHCII from WT, LHCII-qH(−) and LHCII-qH(+). **(B)** (Top panel) Maximum fluorescence emission intensity relative to WT normalized for concentration from steady-state fluorescence emission experiments (squares). Amplitude of the quenched lifetime component from 1D-ILT analysis of the ensemble fluorescence lifetime for LHCII trimers from WT and the LHCII-qH(−) and LHCII-qH(+) mutants (triangles). (Bottom panel) Lifetime components from 1D-ILT fits. **(C)** Schematic of single-molecule fluorescence experiment. 610 nm excitation light is focused on single LHCII immobilized on a coverslip. Green ovals represent visible or scanned LHCII. Gray ovals represent quenched or dark-state LHCII that were not visible during the scan. An example single-molecule intensity trace is shown binned at 20 ms (blue). States were determined using a change-point finding algorithm. The intensity of these states is shown in black (left y-axis) and the lifetime is shown in grey (right y-axis). **(D)** 2D density plots of fluorescence intensity and lifetime for all single-molecule states from LHCII-qH(−) and LHCII-qH(+) samples. Lifetime and intensity states come from individual traces, like those shown in (C).

To investigate the two components present in the ensemble fluorescence, single-molecule fluorescence spectroscopy was performed to uncover the underlying distribution of photophysical behavior in the LHCII-qH(+) and LHCII-qH(−). Each single-molecule time trace shows the fluorescence emission from an individual LHCII. Representative traces are shown in Figure 3C and Figure S9, and representative scans in Figure S10. Within each trace, time periods of emission at a constant intensity were observed with transitions between these periods. The fluorescence lifetime was determined for each period, such that a state of LHCII during that period can be defined by its intensity and lifetime (Figure 3C). The duration of each state is known as the dwell time.

Histograms of the lifetime, intensity, and dwell time from the single-molecule experiments on each sample can be found in Figures S11-S14. Consistent with the ensemble experiments, the median fluorescence intensity of LHCII-qH(+) at the single-molecule level is less than the fluorescence intensity of LHCII-qH(−) (50 counts per 20 ms vs 67 counts per 20 ms, Figures S12-S13). We found that the median fluorescence lifetime in single-molecule experiments of LHCII-qH(+) is shorter than that of LHCII-qH(−) or WT LHCII (1.46 ns in qH(+) vs 1.99 in qH(−) and 1.92 in WT Figures S11-S13) which is consistent with an energy dissipation pathway being promoted by qH. For both samples, the single-molecule lifetimes are shorter than the corresponding ensemble lifetimes (2.32 ± 0.2 ns in qH(+) and 3.43 ± 0.04 ns in qH(−)), likely due to a small degree of photodegradation. The single-molecule lifetime distribution is also limited by the instrument response function of ∼0.4 ns, which buries the short 0.2 ns lifetime component observed in ensemble measurements. The ensemble detector, which was used to measure the amplitude and lifetime of the quenched component, has an instrument response function of less than 0.1 ns and thus did not have the same limitation.

To determine the distribution of states of LHCII, two-dimensional density plots of intensity and lifetime were constructed from all the measured intensity and lifetime values from single-molecule traces (Figure 3D). These plots showed that LHCII-qH(+) and LHCII-qH(−) exhibit a similar photophysical state profile, but that LHCII-qH(+) displays a higher probability of being in lower-intensity, shorter-lifetime states. These data support the hypothesis that the quenching measured in ensemble data of LHCII-qH(+) arises from a quenched subpopulation, rather than a homogenous change in all isolated trimers. Such a subpopulation likely also exists in the LHCII from unquenched samples, as LHCII-qH(−) access the lower-intensity and shorter-lifetime states as well, albeit with a decreased propensity (Figure 3D).

### LHCII-qH(+) show greater state-switching

The dwell times and transitions between states can be used to evaluate the dynamics and connectivity associated with qH. Changes in the emissive state of LHCII arise from conformational changes that impact the emission of the embedded chlorophyll (*38*, *67*). Dwell time is inversely related to state-switching; the dwell time decreases as the frequency of switching between states increases. First, the dwell time histograms were compared for LHCII-qH(+) and LHCII-qH(−). The median was shorter for LHCII-qH(+) trimers than for LHCII-qH(−) trimers at 0.38 seconds and 1.2 seconds, respectively (Figures S12-S13), indicating that LHCII-qH(+) exhibited more switching between states. (*37*, *67*)

Examination of the dwell times for both samples also revealed three patterns, reflecting distinct state-switching behavior: static, dynamic, and blinking (Figure 4A). Static complexes did not change intensity over the course of the approximately 30 second duration of the measurement, dynamic complexes changed intensity but remained in emissive, or “on,” states, and blinking complexes transiently entered non-emissive, or “off,” states. “On” states are defined as states with fluorescence emission intensity above background while “off” states are defined as states with fluorescence intensity below background. All samples contained approximately the same percentage of static complexes (15-20%). The LHCII-qH(−) sample contained a greater percentage of dynamic complexes (53%) as compared to the LHCII-qH(+) sample (36%). In contrast, the LHCII-qH(+) sample contained a greater percentage of blinking complexes (49%) as compared to the LHCII-qH(−) one (∼30%). Notably, almost half the complexes are therefore classified as blinking for the LHCII-qH(+) (Figure 4B). Within blinking complexes, the percentage of time spent in the off state are comparable across samples (Figure S15; ∼22% on average).

**Fig. 4.**
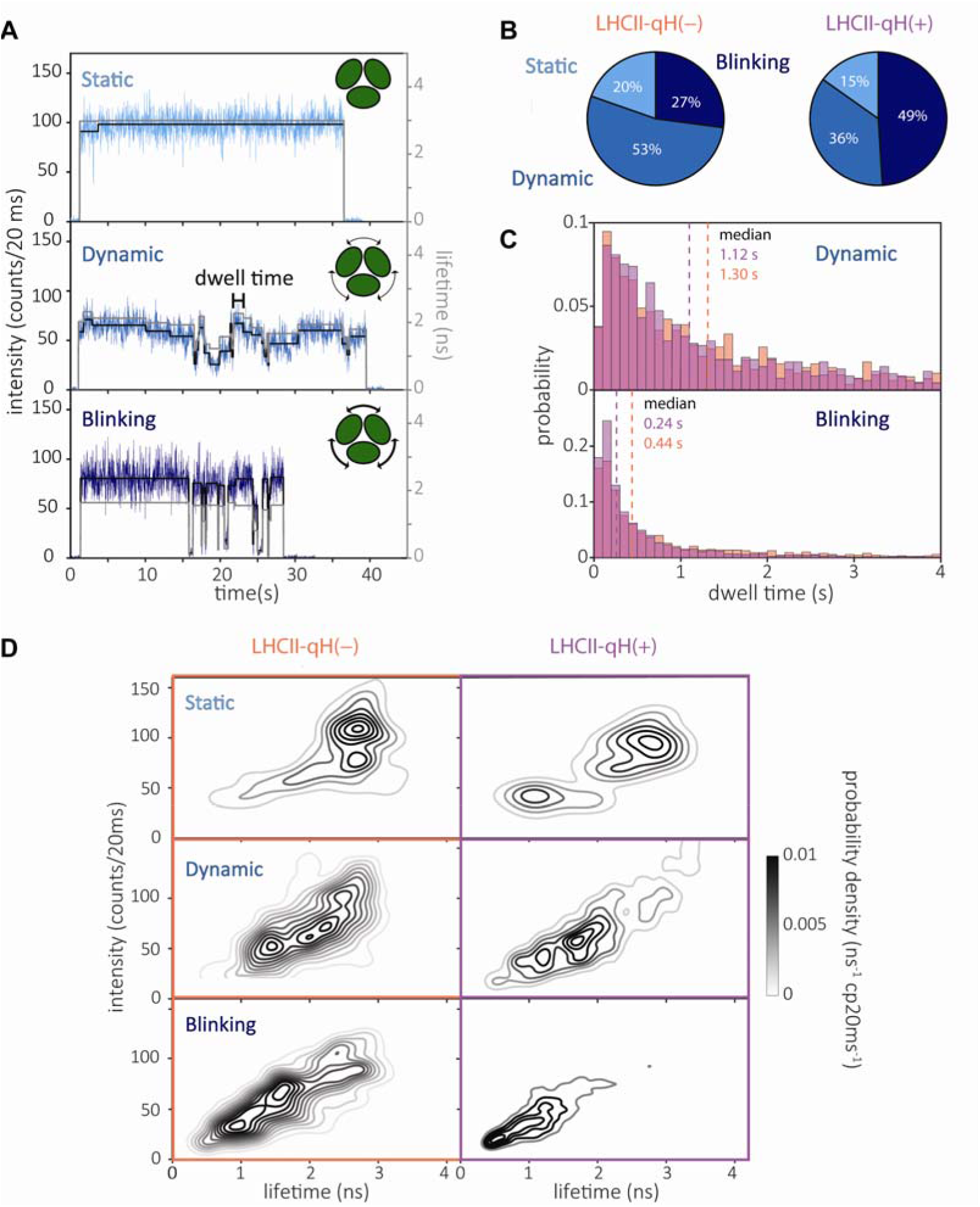
qH is due to a subpopulation of LHCII proteins with enhanced quenching. **(A)** Representative intensity and lifetime traces of the static, dynamic and blinking subpopulations of LHCII. Insets represent state switching (arrows) within LHCII trimers (green). Thicker arrows indicate greater switching. **(B)** Percentage of complexes in each subpopulation from both mutants. **(C)** Histograms of dwell times for dynamic and blinking subpopulations in LHCII-qH(−) and LHCII-qH(+). Dashed lines show the median for each distribution. **(D)** Density plots for LHCII-qH(−) and LHCII-qH(+) by molecule type

To characterize differences in the switching kinetics, the median dwell time was calculated for each type of state-switching within each pattern of behavior. For both LHCII-qH(+) and LHCII-qH(−), the static and dynamic complexes showed similar overall median dwell times at ∼12 seconds for static complexes, ∼1 second for dynamic complexes, and ∼0.3 seconds for blinking complexes (Figure 4C, Figure S15). Blinking molecules have the greatest relative difference in dwell time between the two mutants. The difference in the distribution of states (Figure 4D) suggests that the nature of the transitions changed for the samples. Specifically, analysis of the on-to-on transition for blinking complexes showed a median timescale approximately twice as fast for the LHCII-qH(+) sample (0.24-0.28 s) as for LHCII-qH(−) (0.42-0.54 s) (Figure S15). The difference in the on-to-on switching rate between the two samples could suggest that the on state in the blinking populations varies in its conformational flexibility between LHCII-qH(+) and LHCII-qH(−).

Blinking complexes in both samples are associated with decreased intensity and lifetime (Figure 4D, lower panel ‘Blinking’). However, in the LHCII-qH(+) the decrease is more pronounced: the blinking complexes do not access high intensity and lifetime states. Thus, the LHCII-qH(+) complexes not only contain a larger subpopulation of LHCII trimers undergoing blinking (49% vs 30%), but also the blinking complexes themselves displayed enhanced quenching, suggesting differences in both the population composition and nature of the quenched complexes.

## DISCUSSION

The purification of natively quenched LHCII trimers, made possible by the slow relaxation kinetics of qH, offers a unique opportunity to study antenna-dependent photoprotection in vitro without relying on an artificially-induced system. Here, we thus used the *soq1 roqh1* double mutant, in which qH is constitutively active (*52*) (Figure 1), compared to the *soq1 roqh1 lcnp* triple mutant, in which qH is inactive, to gain insights into the biophysical processes underlying this sustained quenching pathway in LHCII trimers.

Interestingly, we found that qH has a photoprotective effect on LHCII itself, by slowing down the photobleaching process in a suspension of LHCII trimers exposed to high light (Figure 1). Photobleaching has been previously observed in LHCII in detergent environment, as well as in reconstituted lipid membranes, and has been ascribed to the reaction between chlorophyll triplet states in LHCII with O_2_ forming singlet oxygen (*68*). Together with our previous finding that qH prevents lipid peroxidation (*50*), this result further supports the photoprotective role of this NPQ component.

Ensemble and single-molecule fluorescence lifetime analyses show that LHCII trimers with or without qH access a partially overlapping range of fluorescence intensity and lifetime states, but with different probabilities (Figure 3). Surprisingly, blinking complexes in the LHCII-qH(−) sample accessed fluorescence lifetimes and intensities as low as those observed in LHCII-qH(+). However, it remains unclear whether these states arise from the same conformational pathways. The inability of blinking LHCII-qH(+) complexes to access the highest intensity and lifetime states observed in LHCII-qH(<u>−</u>) suggests that qH may alter the conformational landscape of a subset of LHCII trimers, favoring strongly quenched states. Previous single-molecule measurements of LHCII similarly observed an increase in the population of trimers with lower intensity and/or shorter lifetime populations under qE conditions (*12*, *69*). Whether the quenched conformation could be the same one observed in other NPQ components, such as qE, will likely remain an open question until the changes occurring in both qE and qH have been identified. We can however bring here the beginning of an answer.

Blinking, or fluorescence intermittency in single-molecule experiments, occurs when a fluorophore transitions from an emissive “on” state to a non-emissive “off” state, in a reversible manner. If the fluorophore switches irreversibly to a permanent off state, the process is instead referred to as bleaching, equivalent to photodamage of the antenna proteins. Blinking is thought to occur when excitation energy becomes temporarily trapped in a non-emissive electronic state; the precise mechanisms remain incompletely understood (*70*). Previous experiments have attributed LHCII fluorescence intermittency to conformational changes in the protein (*71*, *72*). However, these behaviors were reported to be modulated by pH: upon acidification, switching into dimmer states increased while switching back to brighter states was decreased, resulting in net stabilization of the dim states (*69*, *72*). qH is independent of ΔpH (*51*), and pH was kept invariant in the current study (at pH 8.0). Importantly, the LHCII-qH(+) displays the opposite effect to that caused by pH acidification – we observed a greater frequency of state switching in the sample with dimmer states (lower fluorescence lifetime and intensity) (Figure 4). Furthermore, we resolved distinct subpopulations, where only 27% or 49% of LHCII-qH(−) and LHCII-qH(+), respectively, exhibited blinking. In contrast, under qE-related conditions, including acidification and increased photon flux, the switching rate from “on” to “off” states increased across the sample rather than in a specific subpopulation (*72*). Taken together, these differences suggest that the off states observed in qH conditions occur due to either a different underlying conformational change than the ones triggered by acidic pH or a different trigger for the same conformation.

The model of a unique quenched conformation is also hard to reconcile with the action of pigment changes such as zeaxanthin accumulation, a major part of qE (*73*), when qH does not rely on any significant change in pigment content in LHCII (*49–51*) (Figure S2). The idea of a unique quenched conformation was also recently challenged by simulated measurements which led Gray and coworkers to argue that experimental kinetics of LHCII quenching can only be explained by the co-existence of multiple quenching mechanisms (*71*, *74*). While they do not exclude the possibility of a single quenched state accessible by various modifications, our results rather support the occurrence of various quenched conformations in the LHCII trimer, some of them promoted by the different molecular players of NPQ pathways (e.g. PsbS or LCNP).

Establishing the exact nature of the conformational change responsible for qH quenching is challenging in large part because photophysical differences were only observed in a subpopulation of trimers, making structural changes difficult to observe via ensemble methods. Resonance Raman spectroscopy combined with detergent removal experiments established that the quenching is not correlated with aggregation signatures (Figure 2C,D), demonstrating that qH occurs at the single-trimer level. The absence of differences in the Raman spectra between the LHCII-qH(+) and LHCII-qH(−) samples, while CD does identify a possible involvement of neoxanthin and Chl *b* (Figure 2B), might be explained either by the loss in average of a small change in the heterogeneous samples in the Raman spectra, or by a modification that affects the protein backbone and changes pigments spacing without altering the chemistry of the pigments themselves, as would twisting or hydrogen bonding.

While further experiments will have to test these hypotheses, the results obtained in CD can however point to which pigments might be at play in the energy quenching by qH. CD spectroscopy in Georgakopoulou *et al.* showed that the negative peak around 470 nm was present in neoxanthin knockout mutant (*59*), suggesting that the peak in our data (Figure 2B) could be due to changes in neoxanthin. Changes in this peak have been observed in LHCII upon incorporation in nanodiscs (*11*, *60*), which is correlated with qE; however, in those experiments the peak became more negative rather than less negative, as we observe for LHCII-qH(+) compared to LHCII-qH(−). Furthermore, upon nanodiscs incorporation, there is also a peak shift in the 490 nm region, attributed to interactions of Chl *a*612 with lutein, that we do not observe in our results (*11*, *59*). Another possibility, also based on the findings of Georgakopoulou *et al.*, is that the 470 nm peak is due to interruptions in inter-pigment interactions, as the 470 nm peak is seen in monomeric LHCII. Some of the pigments involved could be the Chl *b*608-b609 and *b*606-b607 pairs (*59*, *61*), also present around helix C of the protein in the vicinity of neoxanthin (see Figure 2A). Interestingly, these pigments matter for energy transfer from monomer to monomer (*7*, *75*). Consistent with this picture, the fluorescence level returned to the control level upon dissociation of trimers into monomers by phospholipase treatment (Figure S3). This loss of quenching upon monomerization, however, is not because trimers are required for qH — indeed monomeric antenna proteins can also be a site for qH (*54*) — but rather because monomerization perturbs pigments at the trimer subunit interface, possibly including neoxanthin and surrounding pigments (*76*). Further experiments will be necessary to determine whether these pigments are involved in forming the quenching site associated with qH, or whether their spectral changes are merely a signature of the quenched conformation.

The question remains of how a subpopulation of quenched trimers can induce the strong light limitation observed in the *soq1 roqh1* mutant (*52*). While a hydrophobic mismatch model has been proposed for qH (*77*), qH is retained in detergent-solubilized isolated LHCII under non-stress conditions, indicating that quenching requires neither the membrane environment nor stress-induced changes to lipid bilayer properties. Interestingly, we observed an increase in quenching in aggregated conditions (Figure S5B), suggesting that qH-active LHCII could act as an energy sink in vivo for neighboring trimers via energy transfer as previously proposed (*78*). Such a model could also explain the decrease in qH intensity observed during purification processes from intact leaves to purified LHCII (*49*): while the quenched LHCII are still present in solution, the progressive disruption of antenna-antenna interactions, from tightly packed membranes to micellar suspensions, decreases the amount of excitation energy dissipated.

Altogether, the results demonstrate that qH relies on a small conformational change that we tentatively attribute to the helix C region and monomer-monomer interface, affecting a subpopulation of LHCII trimers. This change stabilizes LHCII in a conformation characterized by a shorter fluorescence lifetime and an increased fraction (by a factor of two) of complexes displaying blinking. The enhanced propensity for fluorescence intermittency appears directly photoprotective for the antennae, linking the microscopic behavior to biological function. The identification of multiple functional conformations of LHCII complexes, with changes in both properties and populations, may be key to understanding how this important complex performs not only the majority of light harvesting but also a wide range of photoprotective processes.

## MATERIALS AND METHODS

### Plant material and growth conditions

The WT and mutant plants used in this study were from *Arabidopsis thaliana* Col-0 background. The *soq1-1 roqh1-1* and *soq1-1 roqh1-1 lcnp-1* mutants, referred to as *soq1 roqh1* and *soq1 roqh1 lcnp* through the article, were generated and characterized in a previous study (*52*). The *soq1-1* and *soq1-1 lcnp-1* mutants (*soq1* and *soq1 lcnp*) are also described in previous articles (*50*, *51*).

Plant seeds were surface sterilized using 70% ethanol and germinated on agar plates (0.5x Murashige and Skoog Basal Salt Mixture, Duchefa Biochemie, pH 5.7 adjusted with KOH, and agar) after stratification for 24h in the dark at 4°C. The seedlings were grown for 3 weeks under a 12h/12h light/dark regime and 150 μmol photons m^−2^ s^−1^ and 22°C. They were then transferred to soil for five (all lines except *soq1 roqh1*) to seven (*soq1 roqh1*) weeks under short day conditions (8h light/16 h dark) at 150 μmol photons m^−2^ s^−1^ and 22°C/18°C.

### LHCII isolation

Thylakoids were extracted from plants acclimated either to growth light for 1h (WT, *soq1 roqh1*, *soq1 roqh1 lcnp*) or to a 6h cold and high light treatment (4°C, 1600 μmol photons m^−2^ s^−1^) (‘cold high-light’ WT, *soq1* and *soq1 lcnp*). The purification was performed following a protocol described previously (*49*, *79*). All steps were performed in the cold. Briefly, leaves were grinded in B1 solution (20 mM tricine–KOH pH 7.8, 400 mM NaCl, 2 mM MgCl_2_, 0.2 mM benzamidine, 1 mM aminocaproic acid, and 0.2 mM PMSF), filtered through four layers of Miracloth, and centrifuged 5 min at 27,000g at 4°C. The pellet was resuspended in 15 ml B2 solution (20 mM tricine–KOH pH 7.8, 150 mM NaCl, and 5 mM MgCl_2_, 0.2 mM benzamidine, 1 mM aminocaproic acid, and 0.2 mM PMSF), deposited on a 1.3 M/1.8 M sucrose step gradient and ultracentrifuged 30 min in a SW28 rotor at 19,000 rpm and 4°C. The green thylakoid band between both sucrose layers was collected and washed in B3 solution (20 mM tricine–KOH pH 7.8, 15 mM NaCl, and 5 mM MgCl_2_), then centrifuged 15 min at 27,000g and 4°C. Finally, the pelleted membranes were washed in storing solution (20 mM Tris KOH pH 8.0, 0.4 M sucrose, 15 mM NaCl, and 5 mM MgCl_2_), centrifuged 10 min at 27,000g and 4°C, then resuspended in a small amount of storing solution. Chlorophyll concentration was measured by absorption spectroscopy of pigments extracted in acetone according to (*80*). Aliquoted samples were frozen in liquid nitrogen and stored at −80°C until use.

LHCII trimers were isolated by gel filtration chromatography following a protocol described previously (*49*, *79*): a volume of thylakoid membranes corresponding to 400 μg Chl was solubilized at a final concentration of 2 mg Chl mL^−1^ in a final 4% (w/v) n-dodecyl α-D-maltoside (α-DDM) (Anatrace) for 15 min on ice, vortexing briefly every 3 min, then centrifuged at 14,000 rpm for 5 min at 4°C to remove unsolubilized material. Gel filtration was performed on a ÄKTA FPLC chromatography system (Amersham Pharmacia Biotech) coupled to a Superdex 200 Increase 10/300 GL column (GE Healthcare) equilibrated with 20 mM Tris–HCl pH 8.0, 5 mM MgCl_2_, and 0.03% (w/v) α-DDM in the cold. The flow rate was 1 mL min^−1^. The proteins were detected at 280 nm absorbance. Fractions containing LHCII trimers were collected and pooled, and absorption was controlled on a UV-Vis spectrophotometer (UV-2600i, Shimadzu). If not used immediately, aliquots were flash-frozen and kept at -80°C.

### Ensemble steady-state fluorescence measurements

Fluorescence spectra of purified LHCII were taken on a Fluoromax+ spectrofluorometer (Horiba Scientific). The samples were diluted to optical density (OD)_676nm_ ∼ 0.08 in 20 mM Tris-HCl pH 8.0 and 0.03% α-DDM. Room temperature fluorescence emission spectra were recorded from 650 nm to 800 nm with a 1 nm increment. Excitation was set at 625 nm with a 2 nm bandpass for excitation slit and 3 nm for the emission slit. The spectra were normalized on the maximum sample absorption in the Qy peak (as measured in a UV-Vis UV-2600i spectrophotometer, Shimadzu). For 77K fluorescence emission, samples prepared in the same way with the addition of 20% glycerol were measured in liquid nitrogen from 600 to 800 nm, with an excitation at 475 nm, a 2 nm bandpass for excitation and 1 nm for emission.

### Photobleaching

Purified LHCII at an OD_676nm_ ∼ 1 in 20 mM Tris-HCl pH 8.0 and 0.03% α-DDM was measured for photobleaching. The samples were placed in a 96-well plate and exposed to red light at ∼1200 μmol photons m^−2^ s^−1^ for 15 minutes, at room temperature, in a SpeedZen II chlorophyll fluorescence video imager (JBeamBio) (*81*) for fluorescence emission measurement >680 nm (high pass red filter) with saturating pulses, after 30 sec and then every 60 s, of 20 ms duration and excitation of detection at 470 nm.

## HPLC

LHCII trimers were diluted to an OD of ∼ 0.5 in a solution of 80% acetone (final concentration) and 10 mM Tris KOH pH 7.5 and centrifuged at maximum speed to remove protein debris. Pigment analysis was then performed on a C18 column according to a protocol described previously (*82*), with one modification, the pH of solution A maintained to 7.5. Data was plotted and subjected to a two-way ANOVA test using the GraphPad Prism 10.0 software.

### Circular dichroism spectroscopy

The intrinsic circular dichroism (CD) spectra of LHCII trimers were collected using a Chirascan CD spectrometer (software version 4.5.1825.0) at 25.5□°C. Protein solutions of *soq1 roqh1* (LHCII-qH(+)) and *soq1 roqh1 lcnp* (LHCII-qH(−)) were prepared in 10 mM HEPES buffer at pH□7.6, supplemented with 0.04% (w/v) α-DDM. The samples were at OD of approximately 0.8 at the Qy absorption maximum and measured in a quartz cuvette with a pathlength of 1□mm. The instrument was continuously purged with nitrogen gas to eliminate atmospheric oxygen and prevent ozone formation. The background spectrum was recorded using a solution of the same composition as the sample but without the protein to account for any buffer contributions. Measurements were performed as follows: for each sample, three runs were accumulated and recorded over the wavelength range of 400□nm to 750□nm, with a time-per-point of 1□s and a bandwidth of 1□nm. Data processing and analysis were conducted using OriginPro 2023 version 10.0 software.

### Monomerization of LHCII trimers

300 μg mL^−1^ of LHCII trimers purified as described above were broken into monomers by a treatment with 10 μg mL^−1^ of phospholipase A2 from honeybee venom (Sigma Aldrich) in a buffer containing 20 mM Tris HCl pH 8.0, 0.01% α-DDM and 20 mM CaCl_2_. The samples were incubated for 22h at 22°C with agitation, then deposited on sucrose gradients prepared by freezing and thawing of a solution containing 20 mM Tris HCl pH 8.0, 0.35 M sucrose and 0.03% α-DDM. Separation was done by ultracentrifugation in a SW55 rotor (Beckman Coulter) running at 55,000 RPM for 3h. Monomerized trimers and leftover trimers were taken from the gradient and absorption and fluorescence measured as described above.

### Resonance Raman vibrational spectroscopy

LHCII trimers were purified as described above, using n-dodecyl β-D-maltoside (β-DDM) (Anatrace) instead of α-DDM, as the specific resonance markers of LHCII aggregation are more visible in β-DDM than they are in α-DDM (*22*) and qH-LHCII trimers displayed the same fluorescence behavior after purification in β-DDM under the tested experimental conditions (Figure S16).

Resonance Raman spectra were obtained at 77 K in a liquid nitrogen flow cryostat (Air Liquide), using a Jobin-Yvon U1000 Raman spectrophotometer equipped with a liquid-nitrogen-cooled charge-coupled-device detector (Spectrum One, Jobin-Yvon). Laser excitations at 488.0 nm and 441.6 nm were obtained with Coherent argon (Sabre) and a Liconix helium-cadmium lasers, respectively.

LHCII quenched by aggregation was prepared by detergent removal using SM-2 bioabsorbent beads (Biorad) (see Aggregation tests section), allowing for a 8-fold reduction in fluorescence yield as determined by a mini-PAM-I fluorimeter (Heinz Walz).

### Aggregation tests

250 μL of LHCII trimers, from WT, *soq1 roqh1* or *soq1 roqh1 lcnp*, at OD_676nm_ ∼ 1 were incubated with 2-3 mg of Bio-Beads SM-2 (Bio-Rad) for 2 h in the dark at room temperature with agitation to remove detergent. They were then centrifuged for a few seconds in a bench minicentrifuge to sediment the beads, after which the supernatant was transferred in another tube and immediately measured for steady-state room temperature fluorescence emission as described above, except for sample dilution, which was done in 20 mM Tris-HCl pH 8.0 without detergent.

For tests in various detergent concentrations, purified LHCII samples were incubated in 20 mM Tris-HCl pH 8.0 with α-DDM concentration adjusted to 0.03%, 0.1% or 1%, respectively. Steady-state room temperature fluorescence was then measured as described.

### Ensemble fluorescence lifetime measurements

Fluorescence lifetimes were determined by time-correlated single photon counting (TCSPC) and fit with both least-squares exponential decay fitting and with model-free one-dimensional inverse Laplace transform (1D-ILT) analysis. Broadband excitation was created by coupling the emission of a Ti-sapphire oscillator (MaiTai BB, SpectraPhysics; centered at 800 nm, 80 MHz repetition rate) into a nonlinear photonic crystal fiber (FemtoWhite 800, NKT Photonics). A 630–655 nm bandpass filter (ET645/30x; Chroma) was used to select the excitation wavelength. The excitation was focused on a 10 mm x 2 mm quartz cuvette (Helma Analytics) with an average power of 75 nW. Emission from the sample was passed through a 664–715 nm bandpass filter (ET690/120x, Chroma) and detected with a single photon avalanche photodiode (PDM Series, Micro Photon Devices). Photon arrival time was measured with a TCSPC module (PicoHarp 300, PicoQuant, Inc.). The instrument response function (IRF) was measured by collecting scattered light without an emission filter from a 1:200 v:v mixture of HS-40 colloidal silica (Sigma-Aldrich) and water. The full width half max (FWHM) of the IRF was checked each day that data was collected and was 80-90 ps. Data was collected in two ways. Data collected in a pre-binned format was convoluted with the IRF and fit to a biexponential decay using least squares fitting. Data binning was determined by the Picoharp software. Each measurement was performed until at least one million photons were collected and repeated two or three times for technical replicates. Quasi-biological replicates were performed on samples isolated in the same protein preparations but frozen in separate aliquots. Three technical replicates of data collected in time-tagged time-resolved mode for 10 minutes were performed on these quasi-biological replicates. Data collected for 1D-ILT analysis was collected in a time-tagged time-resolved mode for at least ten minutes. Three technical replicates were performed for each sample.

## 1D-ILT Fit to time-tagged time-resolved data

1D-ILT was performed to extract the number of lifetime components in a model-free manner. In short, the photons were transformed from the time domain to the lifetime domain. This was done using the maximum entropy method as described previously (*83*–*85*). The amplitude a_1,2_ and lifetime τ_1,2_ and of each component were determined by:

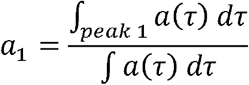

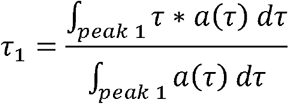

Error bars on the amplitude and lifetime were determined by running the analysis on three technical replicates of the ensemble lifetime measurement and taking the standard deviation of the measurement.

### Single-molecule fluorescence experiment

LHCII trimers were diluted to approximately 40 pM in 20 mM Tris HCl pH 8.0, 5 mM MgCl_2_, 0.03% α-DDM and 10% glycerol with 1% w/v polyvinyl alcohol for immobilization of the protein. To prevent oxidation an oxygen scavenger was used at a final concentration of 2.5 mM protocatechuic acid (PCA) and 25 nM protocatechuate-3,4-dioxygenase (*86*) and single-molecule measurements were performed under a light flow of argon. 80 µL of sample was spin coated onto a glass coverslip using an RPM of 3000 for 30 seconds. Prior to spin-coating glass coverslips were cleaned by sonication in methanol (30 minutes), double deionized water (five minutes), 1M potassium hydroxide (30 minutes), double deionized water (30 minutes). Empty coverslips were stored at room temperature in double deionized water and sonicated for five minutes immediately prior to spin coating. All solutions for coverslip cleaning were vacuum filtered with 200 µm bottle filters.

Single-molecule measurements were performed on a lab-built confocal microscope described previously (*38*). Excitation was performed at 610 nm by a tunable fiber laser (FemtoFiber pro, Toptica; 130 fs pulse duration, 80 MHz repetition rate). Stray light was prevented from entering the microscope using a bandpass excitation filter A630–655 nm (ET645/30x, Chroma). Excitation at 75 nW before the objective was focused using an oil immersion objective (UPLSAPO100XO, Olympus, NA 1.4). Excitation and emission were separated using a dichroic mirror (ZT647rdc, Chroma Technology) and emission was filtered using two bandpass filters (ET700/75m, Chroma and ET690/120x, Chroma) before being detected by an avalanche photodiode (SPCM-AQRH-15, Excelitas). Photon arrival times were recorded using a TCSPC module (TimeTagger20, Swabian Instruments). The IRF was measured on each day of measurements by collecting the scatter off of a coverslip without an emission filter. The IRF was 380-400 ps at FWHM.

During the experiment, a 5 µm x 5 µm area of the coverslip was quickly scanned at an initial rate of 0.05 µm per second using a piezoelectric stage (Mad City Labs, Nano-LP100) to prevent photobleaching. The excitation was then centered at the brightest spot of each molecule and fluorescence was recorded for approximately 30 seconds. Prior to single-molecule experiments the ensemble lifetime of the sample was measured on the confocal microscope using a 20 µL droplet of sample at approximately OD 0.8 per cm to verify sample integrity. At least two single-molecule experiments from the same biological preparation were performed per sample.

### Analysis of single-molecule data

Single-molecule intensity and lifetime state analysis was performed largely as previously described (*38*). Briefly, fluorescence emission was binned at 20 milliseconds to create intensity traces. Changes in intensity were determined using the change point finding algorithm of Watkins and Yang (*87*). Complex behavior was determined by analyzing intensity states. The background is noise in the detector (electronic noise). A background measurement was performed by measuring an empty area of the cover slip on each measurement day. Complexes that did not show more than ten percent deviation in intensity were defined as static. Complexes that showed more than ten percent deviation in intensity but did not display intensity below background were defined as dynamic. Complexes that displayed intensity below background were defined as blinking. Error bars on the percentage of complexes displaying each behavior type were determined by taking the standard deviation of 1000 bootstrap samples.

For each intensity state determined by the intensity change point finding algorithm, the lifetime was fit with a single exponential decay using maximum likelihood estimation (*88*). Two-dimensional kernel density estimation was used to create density plots of fluorescence intensity and lifetime data to better visualize photophysical states (*89*).

## Acknowledgments

We thank Henry Lam for assistance in preparing some of the figures for publication and Stefano Caffarri and Ricardo Javier Vázquez for critical reading of the manuscript.

## Author contributions

Funding acquisition: A.M. and G.S.S.-C. Conceptualization: A.C., M.P.H., G.S.S.-C and A.M. Investigation: A.C., M.P.H., C.I. and E.C.-S. Analysis: A.C., M.P.H., C.I., and E.C.-S. Supervision: A.M., G.S.S.-C, A.P., B.R. and E.R. Writing— original draft: A.C. and M.P.H. Writing—review and editing: A.C., M.P.H., C.I., E.C.-S., A.P., B.R., and G.S.S.-C. and A.M.

## Data, code, and materials availability

All data needed to evaluate the conclusions in the manuscript are present in the manuscript or the Supplementary Materials or the Source Data file. Data are available at Zenodo (https://zenodo.org/records/20936059).

