## Supplemental Data for "Sustained photoprotection involves enhanced fluorescence intermittency in a subpopulation of LHCII"

**This PDF file includes:**

Supplementary Methods
Figs. S1 to S16
Tables S1
References (1 to 4)

### Supplementary Methods

***Estimation of Percent Molecules Blinking***

For WT LHCII or each mutant, the total percent time blinking molecules spent in the off state was calculated by adding up the time of all the dark states and dividing by the total time of the blinking molecule traces for that sample. The total percentages were 24% for WT, 20% for LHCII-qH(−) from *soq1 roqh1 lcnp*, and 23% for LHCII-qH(+) from *soq1 roqh1*. This suggests that molecules were visible only 76%, 80%, and 77% of the time. The number of observed blinking molecules were then divided by the percent time of visibility and multiplied by 100 and rounded to the nearest integer to determine the corrected number of blinking molecules. This was added to the number of static and dynamic molecules to determine the total number of molecules for each sample and then percentages could be determined from there.

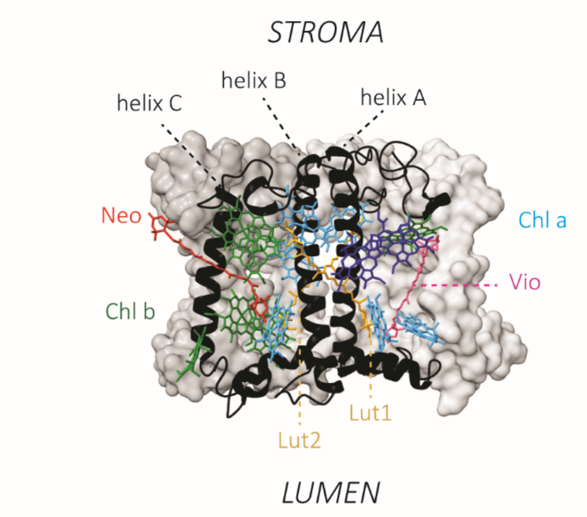


**Figure S1. Structural model of a LHCII trimer in side view, with the structure of one of the monomers detailed and all pigments displayed.**

The protein backbone is displayed in black as ribbon, with the three transmembrane helices indicated. Pigments include chlorophyll *a* (cyan – including the terminal emitter chlorophyll pair in darker blue), chlorophyll *b* (green), and four carotenoids: one violaxanthin (Vio, magenta), two luteins (Lut, orange) and one neoxanthin (Neo, red) are present under non-stress conditions (*1*, *2*). The phytol chains of the chlorophylls are not displayed for clarity. The structure was adapted from 8IWZ (*3*).

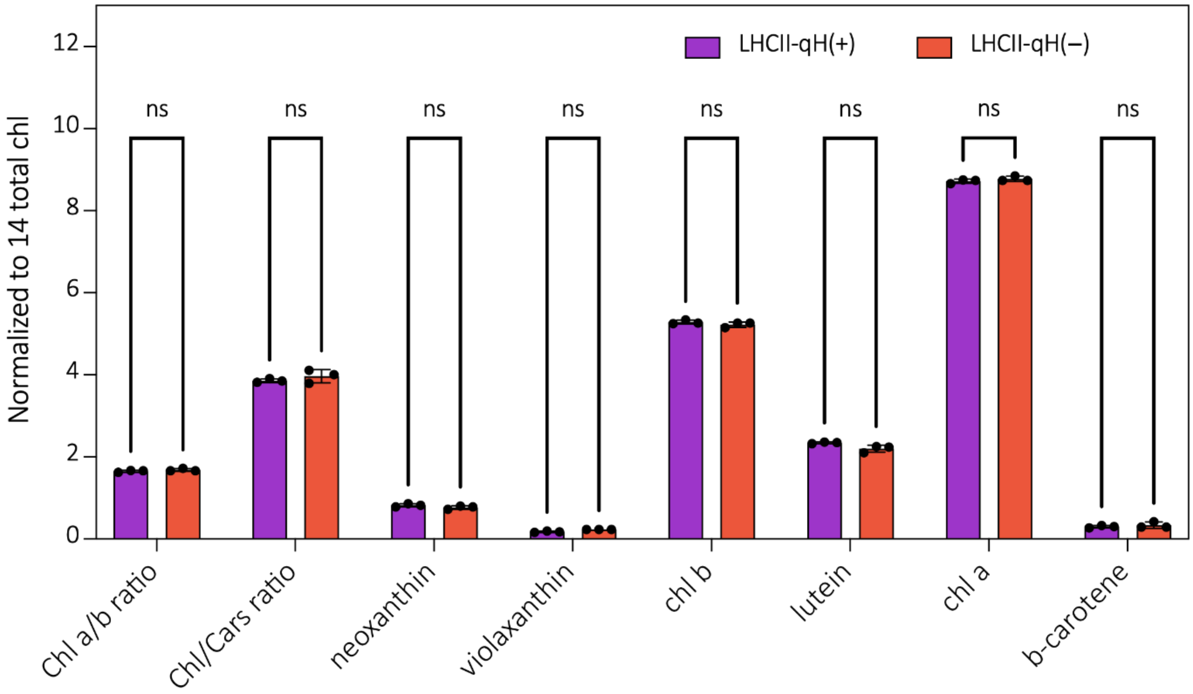


**Figure S2. Pigment content of purified LHCII trimers as determined by HPLC.**

Data represent mean ± SD of three biological replicates, normalized to the 14 Chl content of a single Lhcb monomer. Presence of β-carotene indicates a minor contamination by PSII cores, though similar in both LHCII samples and which therefore cannot explain differences observed between samples. Statistical analyses using a two-way ANOVA test show no significant difference in pigment content of LHCII-qH(−) compared to LHCII-qH(+) ($p$ <0.05).

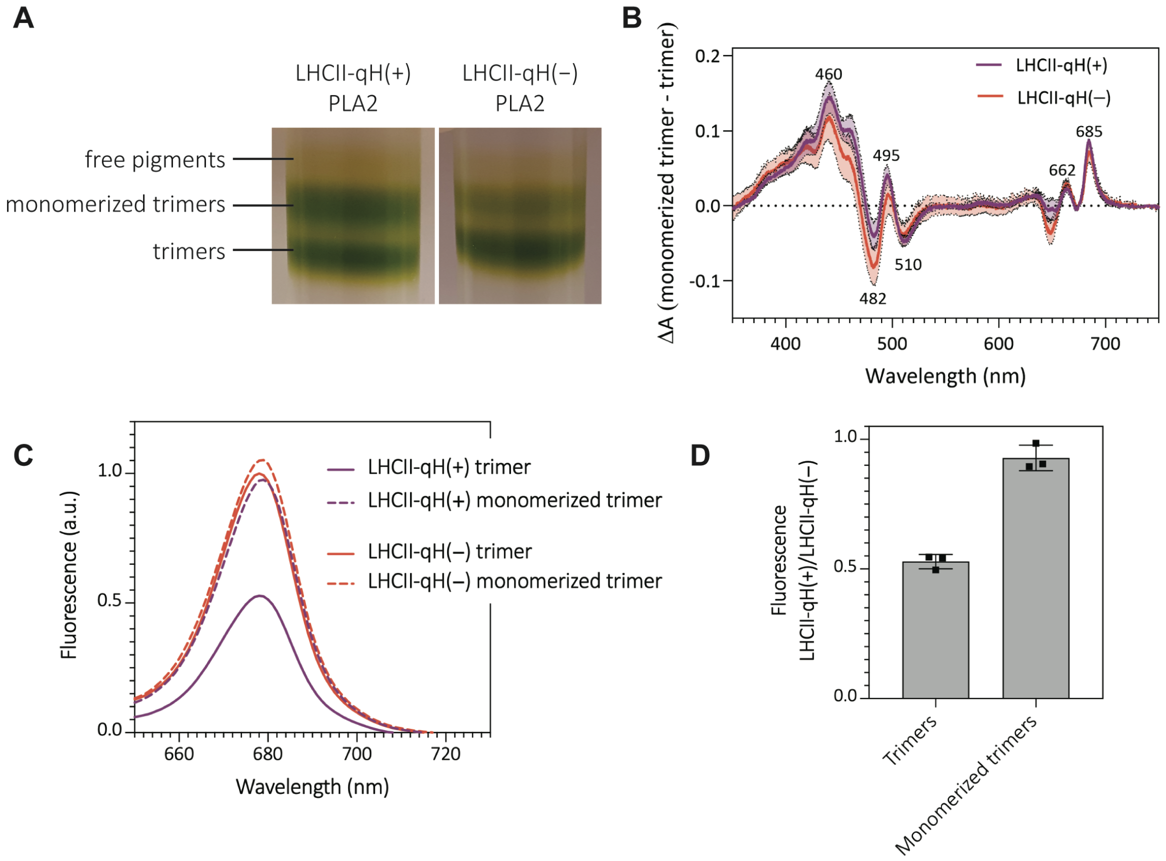


**Figure S3. Phospholipase A2 treatment recovers fluorescence in LHCII-qH(+).**

**(A)** Purified LHCII trimers were treated with phospholipase A2 (PLA2) to separate them into monomers (*4*). **(B)** The absorption difference between the monomerized trimers and the remaining intact trimers is similar for both samples and in line with expected absorption and pigment perturbations after treatment (*4*). **(C)** Room temperature fluorescence emission spectra of isolated LHCII upon excitation at 625 nm and normalized on the maximum sample absorption in the Qy peak. Phospholipase-induced monomerization almost fully recovers the emission of the LHCII-qH(+) sample. (**D)** Fluorescence ratios of the samples presented in C. Data represent mean ± SD of three biological replicates (different thylakoid preparations).

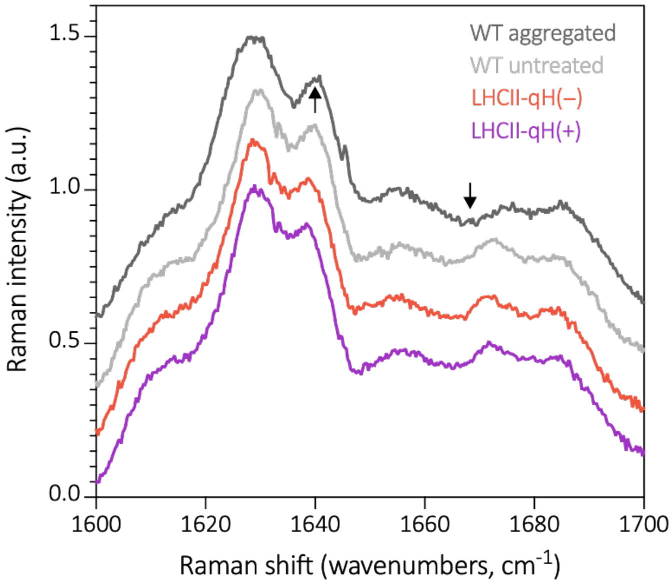


**Figure S4. Resonance Raman spectra with excitation at 441 nm (Chl *b*).**

As observed at 488 nm, the spectra obtained for LHCII-qH(+) (purple) and LHCII-qH(−) (orange) are identical to each other and to the untreated WT (light grey), but they differ from aggregated WT (dark grey), which are quenched to the same extent as LHCII-qH(+) upon detergent removal (arrows indicate the affected regions).

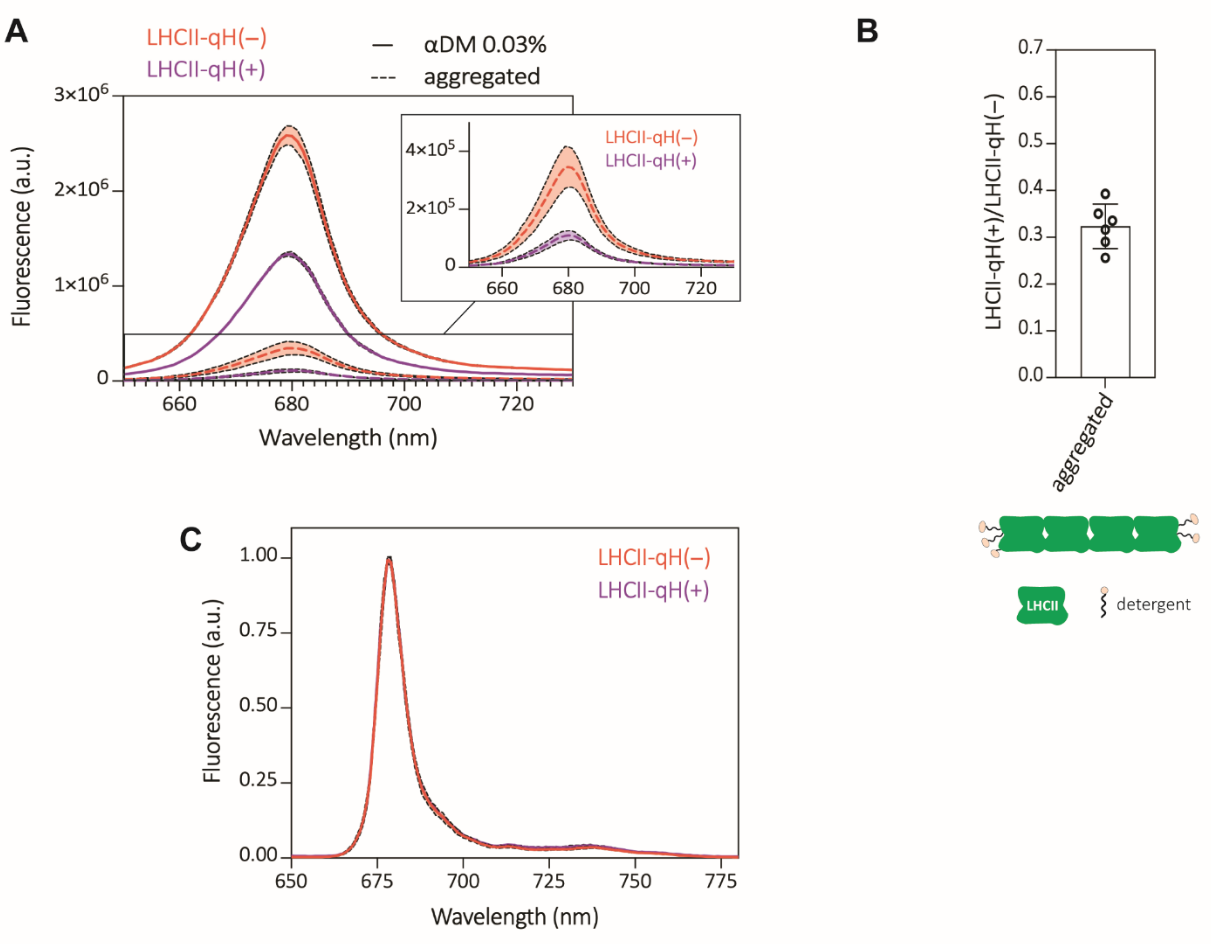


**Figure S5. qH does not rely on LHCII aggregation.**

**(A)** Room temperature fluorescence measurements of isolated LHCII trimers, either aggregated by detergent removal (dashed lines and inset) or solubilized in 0.03% α-DDM detergent. Aggregation decreased the fluorescence of both samples as expected, but without canceling the qH-dependent difference. Data represent mean ± SD (n=4 (for the samples in detergent) and n=6 (for the aggregated samples) replicates (independent LHCII purifications from two different thylakoid batches). **(B)** Ratio of fluorescence of the aggregated LHCII-qH(+) and LHCII-qH(−) samples from panel A. The ratio is lower than the one obtained for trimers in detergent solution, shifting from ~0.5 to ~0.3. i.e. a 70% quenching (Figure 2D). The scheme represents detergent conditions, with LHCII in green and detergent in beige and black. **(C)** 77K fluorescence of purified LHCII trimers, normalized to the maximum of emission.

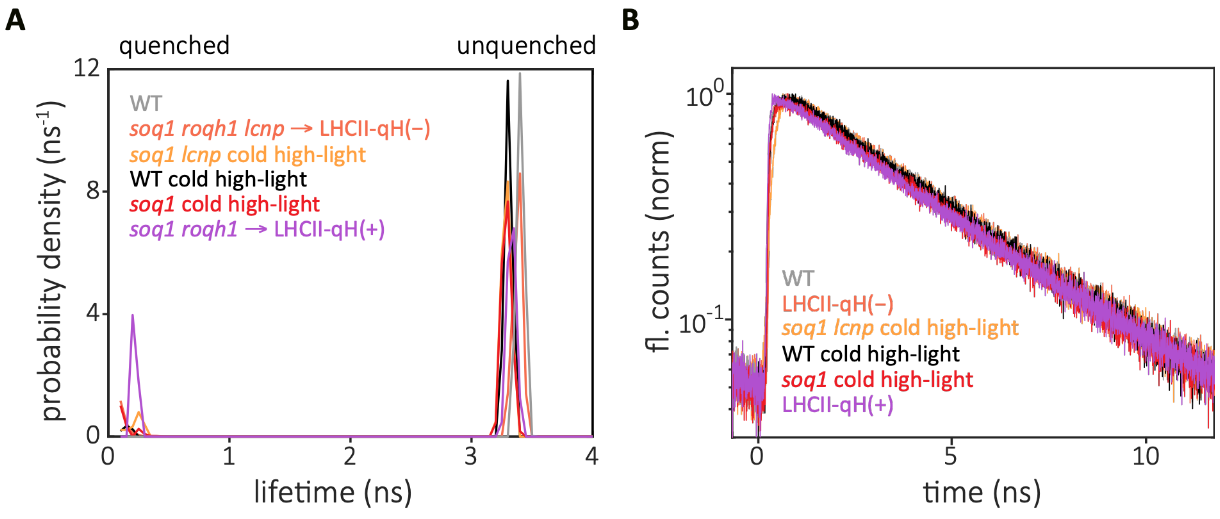


**Figure S6. Fluorescence lifetime measurements for LHCII from all mutants.**

**(A)** Results of 1D-ILT analysis on all samples. **(B)** Fluorescence decay traces from time correlated single photon counting experiments from all samples.

| **A** | | WT | *soq1 lcnp*<br>cold high light | *soq1 roqh1 lcnp*<br>LHCII-qH(−) | WT<br>cold high light | *soq1*<br>cold high light | *soq1 roqh1*<br>LHCII-qH(+) |
|---|---|---|---|---|---|---|---|
| | A1<br>(%) | 7.50<br>(6.88) | 8.54<br>(2.26) | 0<br>(0) | 3.45<br>(5.05) | 6.77<br>(8.54) | 28.66<br>(0.55) |
| | T1<br>(ns) | 0.14<br>(0.04) | 0.23<br>(0.04) | NA | 0.12<br>(0.04) | 0.12<br>(0.03) | 0.22<br>(0.01) |
| | A2<br>(%) | 92.50<br>(6.88) | 91.46<br>(2.26) | 100<br>(0) | 96.55<br>(5.05) | 93.23<br>(8.54) | 71.34<br>(0.55) |
| | T2<br>(ns) | 3.38<br>(0.03) | 3.29<br>(0.01) | 3.40<br>(0.02) | 3.30<br>(0.01) | 3.29<br>(0.02) | 3.33<br>(0.0) |
| | Tavg<br>(ns) | 3.14<br>(0.24) | 3.03<br>(0.08) | 3.40<br>(0.02) | 3.20<br>(0.15) | 3.08<br>(0.25) | 2.44<br>(0.02) |
| **B** | A1<br>(%) | 4.97<br>(0.26) | 10.25<br>(1.93) | 3.73<br>(0.12) | 6.03<br>(1.29) | 19.09<br>(3.91) | 35.9<br>(3.18) |
| | T1<br>(ns) | 0.22<br>(0.01) | 0.21<br>(0.02) | 0.24<br>(0.01) | 0.23<br>(0.02) | 0.21<br>(0.03) | 0.18<br>(0.02) |
| | A2<br>(%) | 95.03<br>(0.26) | 89.75<br>(1.93) | 96.27<br>(0.12) | 93.97<br>(1.29) | 80.92<br>(3.91) | 64.10<br>(3.18) |
| | T2<br>(ns) | 3.48<br>(0.01) | 3.41<br>(0.01) | 3.55<br>(0.03) | 3.44<br>(0.00) | 3.5<br>(0.00) | 3.52<br>(0.02) |
| | Tavg<br>(ns) | 3.32<br>(0.00) | 3.08<br>(0.05) | 3.42<br>(0.03) | 3.25<br>(0.04) | 2.87<br>(0.13) | 2.32<br>(0.10) |

**Table S1. Fluorescence lifetime fitting results for 600 nm excitation.**

**(A)** Results of 1D-ILT analysis on fluorescence decay traces from TCSPC experiments with excitation at 600 nm. Results are averaged across three measurements. Error is standard deviation from these measurements. **(B)** Results of fitting fluorescence decay traces from 600 nm TCSPC experiments with a two-component exponential decay function. Values shown are averages from two quasi-biological replicates (i.e. measures on different aliquots of the same LHCII purification batch), with two to three technical replicates each as shown in Supplemental Figure 7. Error is standard deviation from the two quasi biological replicates.

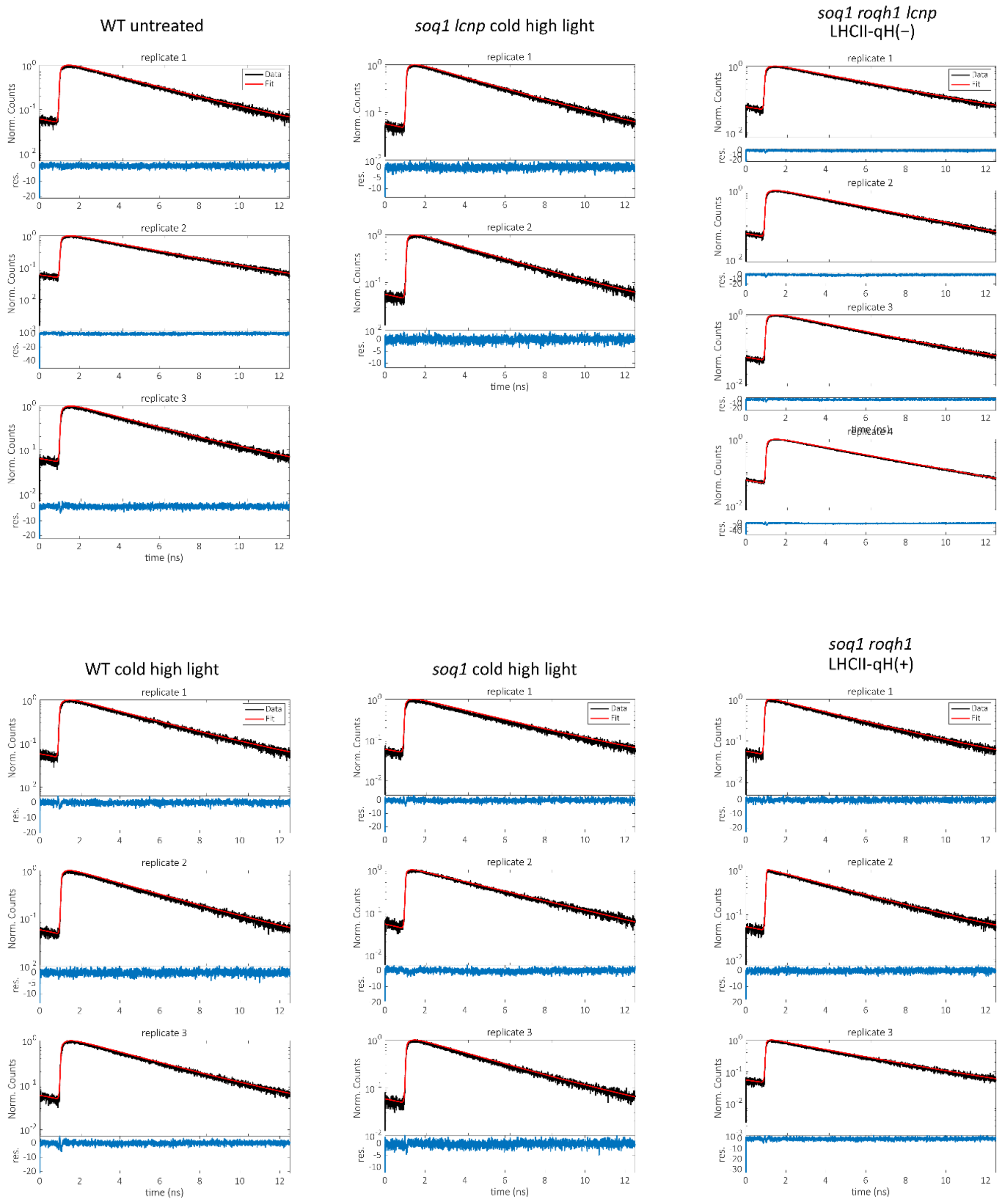


**Figure S7. Two-component least squares ensemble fluorescence lifetime fits.**
Fluorescence decay traces (black) line fit to a two-component exponential decay (red line), with the residual shown below (blue line) for each technical replicate of the first biological replicate for each sample.

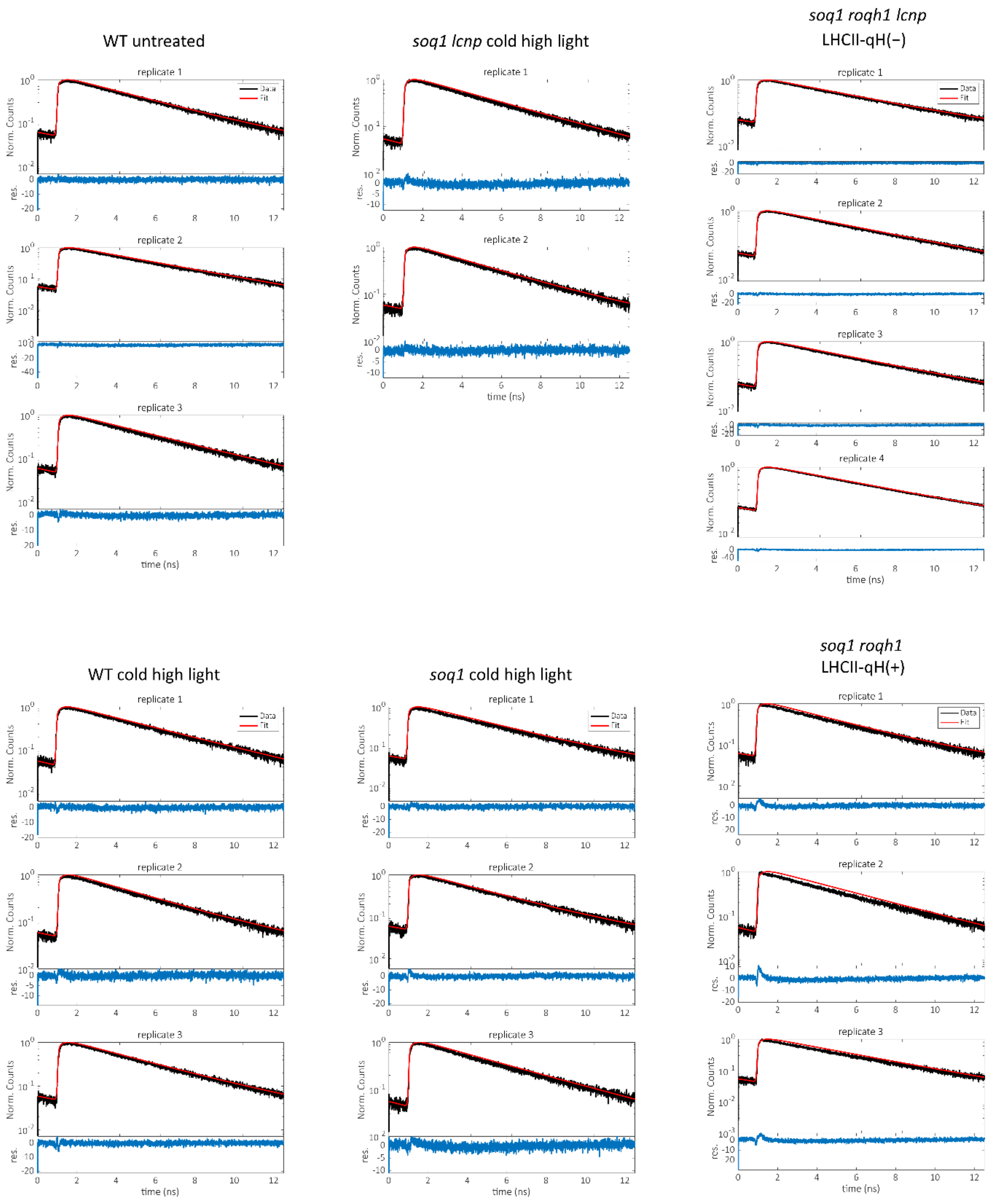


**Figure S8. One-component least squares ensemble fluorescence lifetime fits.** Fluorescence decay traces (black) line fit to a one-component exponential decay (red line), with the residual shown below (blue line) for each technical replicate of the first biological replicate for each sample.

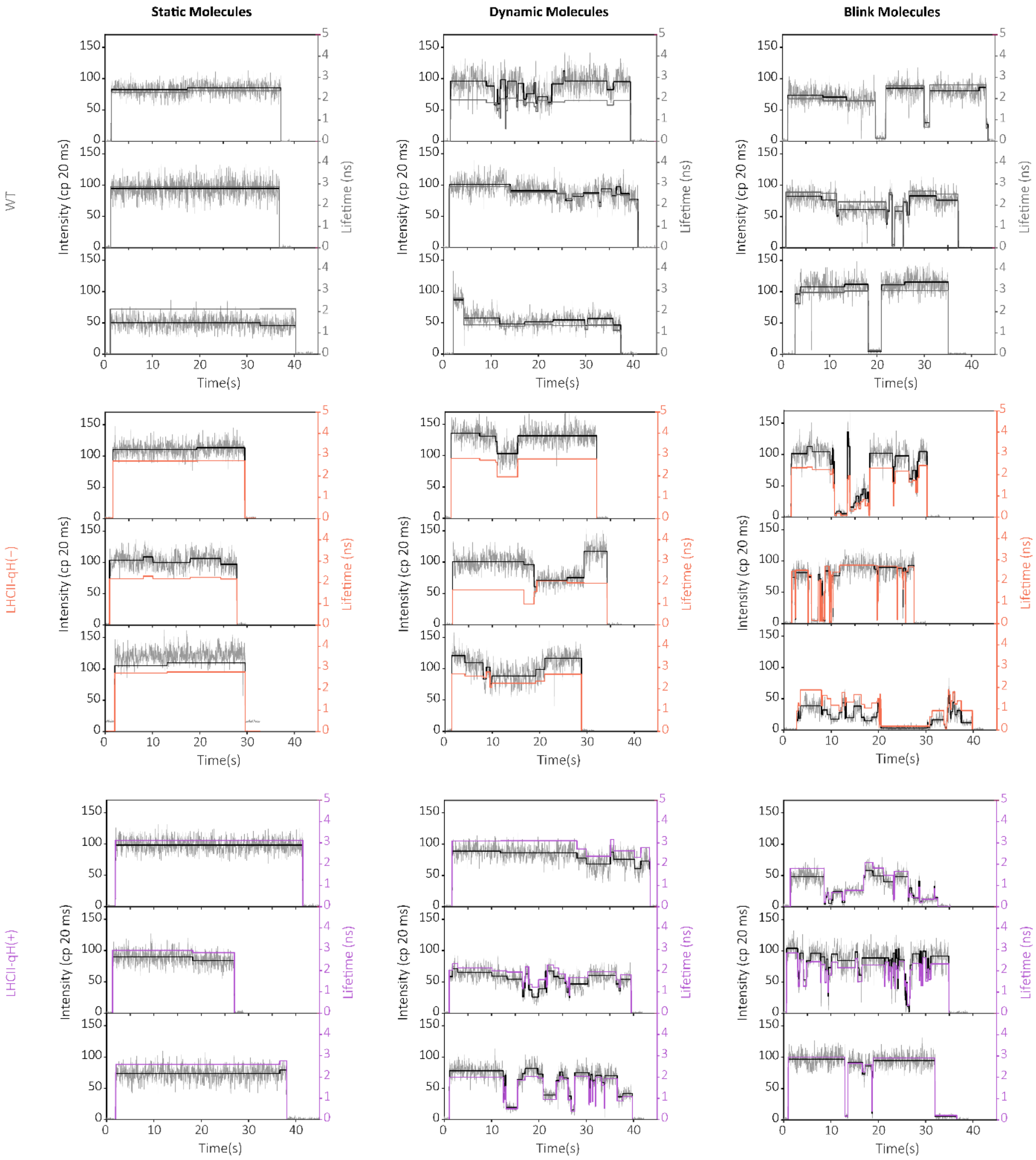


**Figure S9. Example single-molecule traces.**

Example traces of WT (A), LHCII-qH(−) (B) and LHCII-qH(+) (C) and each molecule type.

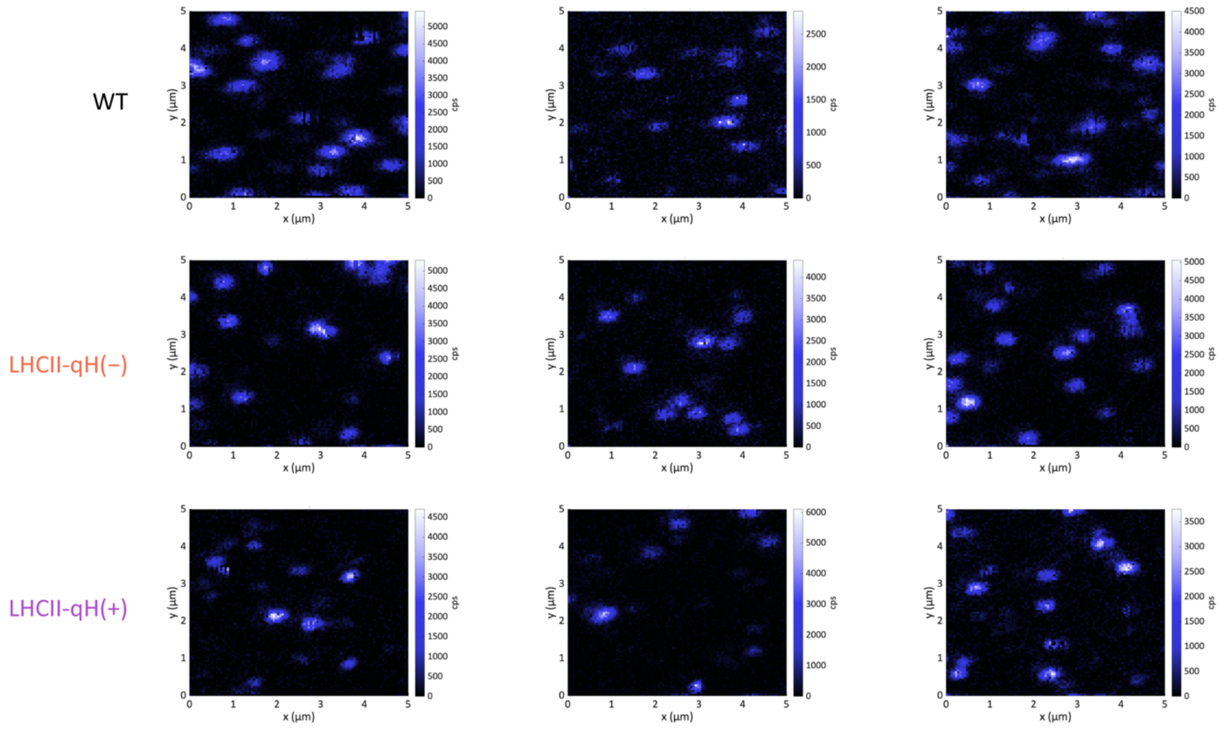


**Figure S10. Example single-molecule scans.**

Example scans from single-molecule experiments with similar concentrations of LHCII deposited on coverslip.

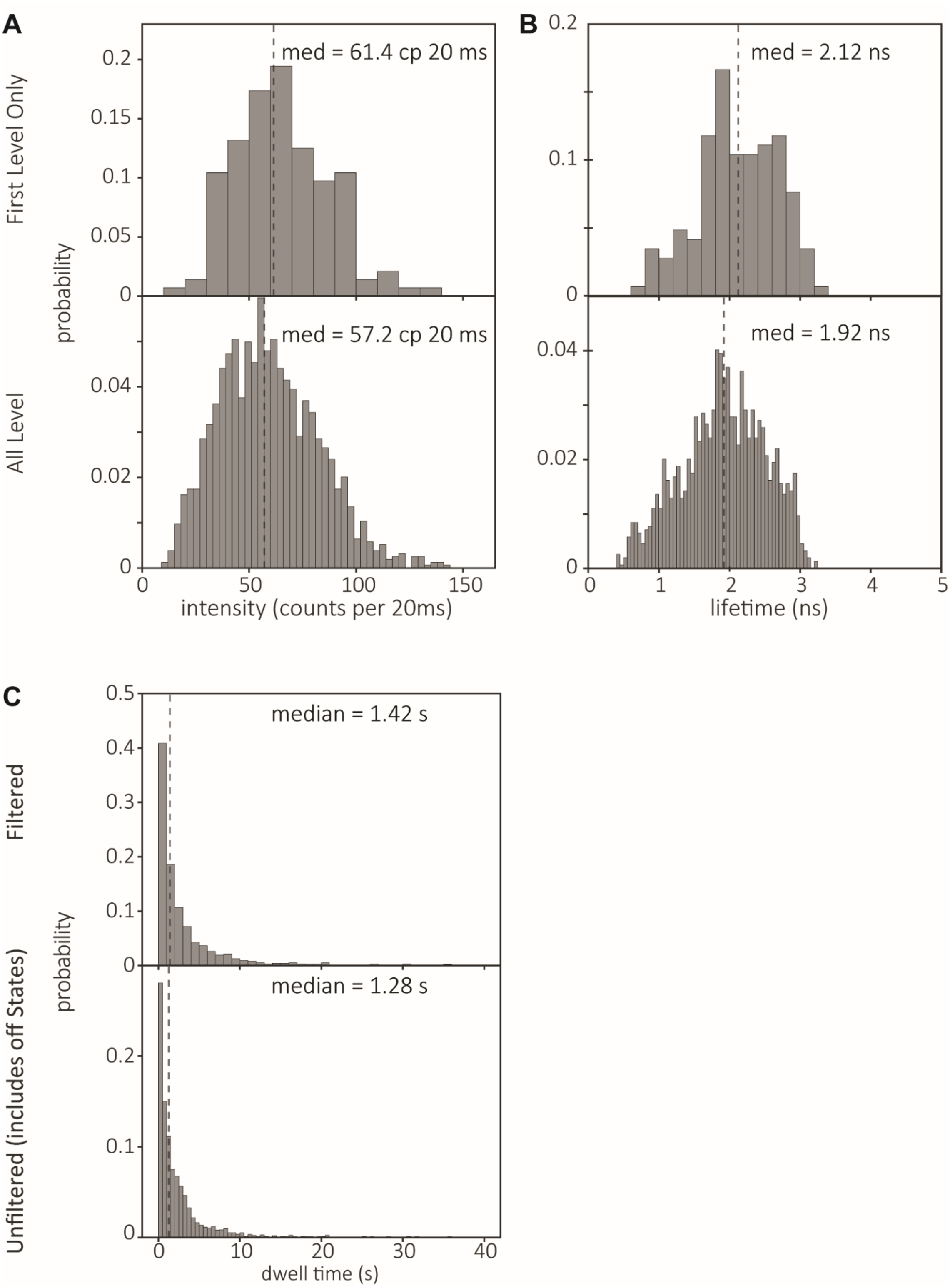


**Figure S11. WT single-molecule data histograms.**

WT, first and all level histograms for intensity (A) and lifetime (B) levels. Median is shown with a dashed line. C. Filtered and unfiltered dwell times. Unfiltered dwell times include the off states that are filtered out in other single-molecule analysis because of limited photon count.

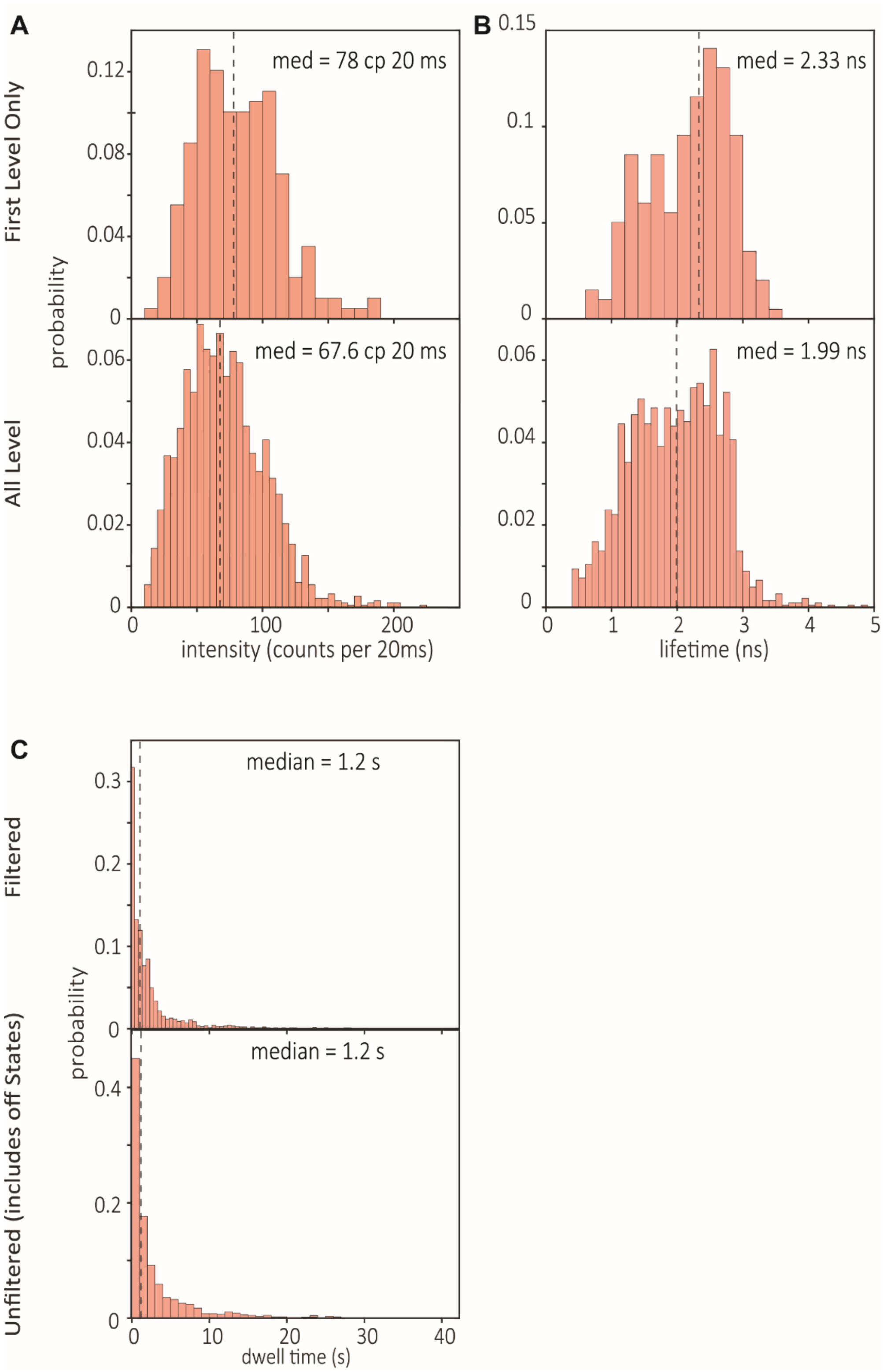


**Figure S12. LHCII-qH(−) single-molecule data histograms.**

LHCII-qH(−) from *soq1 roqh1 lcnp*, first and all level histograms for intensity (A) and lifetime (B) levels. Median is shown with a dashed line. C. Filtered and unfiltered dwell times. Unfiltered dwell times include the off states that are filtered out in other single-molecule analysis because of limited photon count.

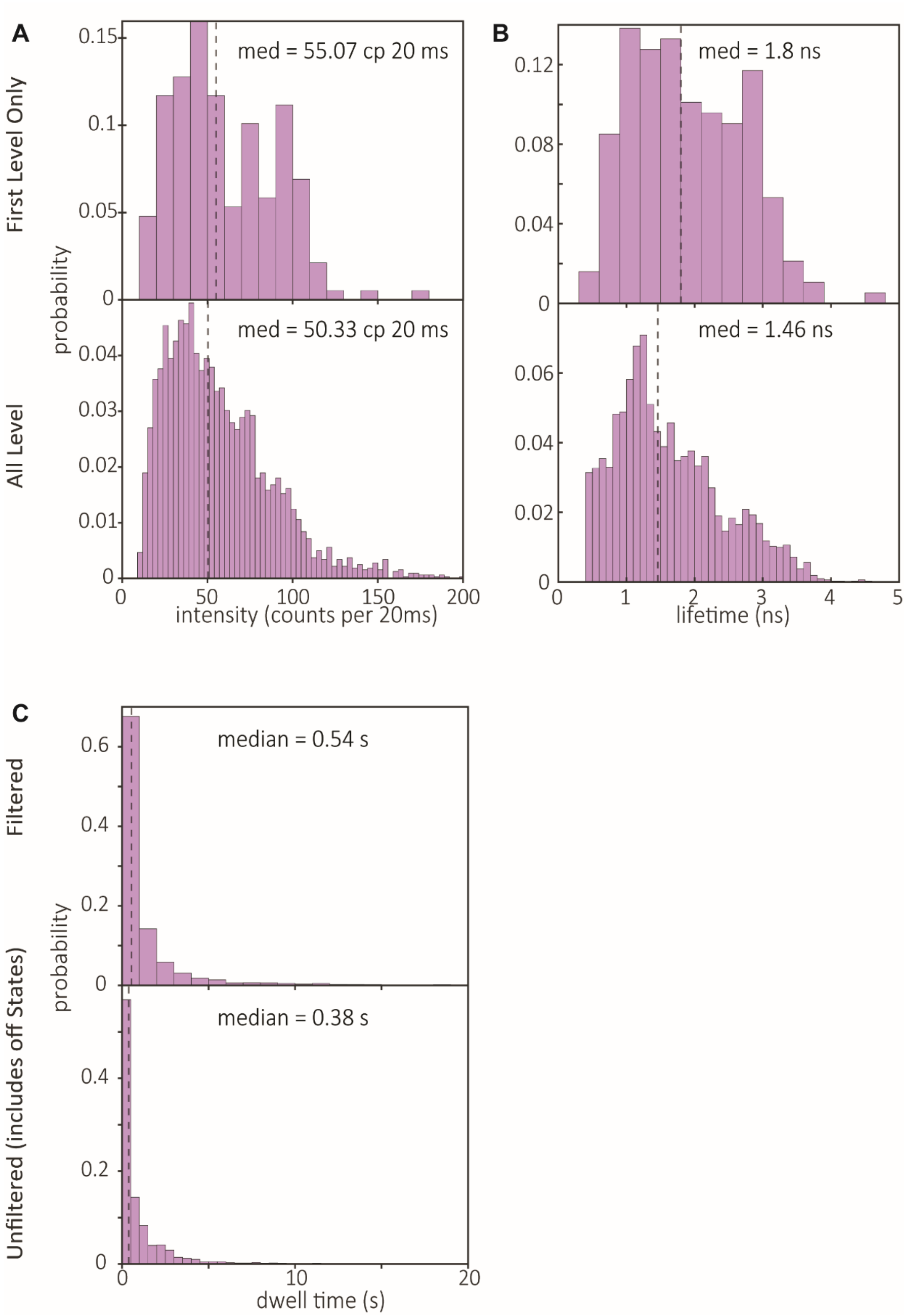


**Figure S13. LHCII-qH(+) single-molecule data histograms.**

LHCII-qH(+) from *soq1 roqh1*, first and all level histograms for intensity (A) and lifetime (B) levels. Median is shown with a dashed line. C. Filtered and unfiltered dwell times. Unfiltered dwell times include the off states that are filtered out in other single-molecule analysis because of limited photon count.

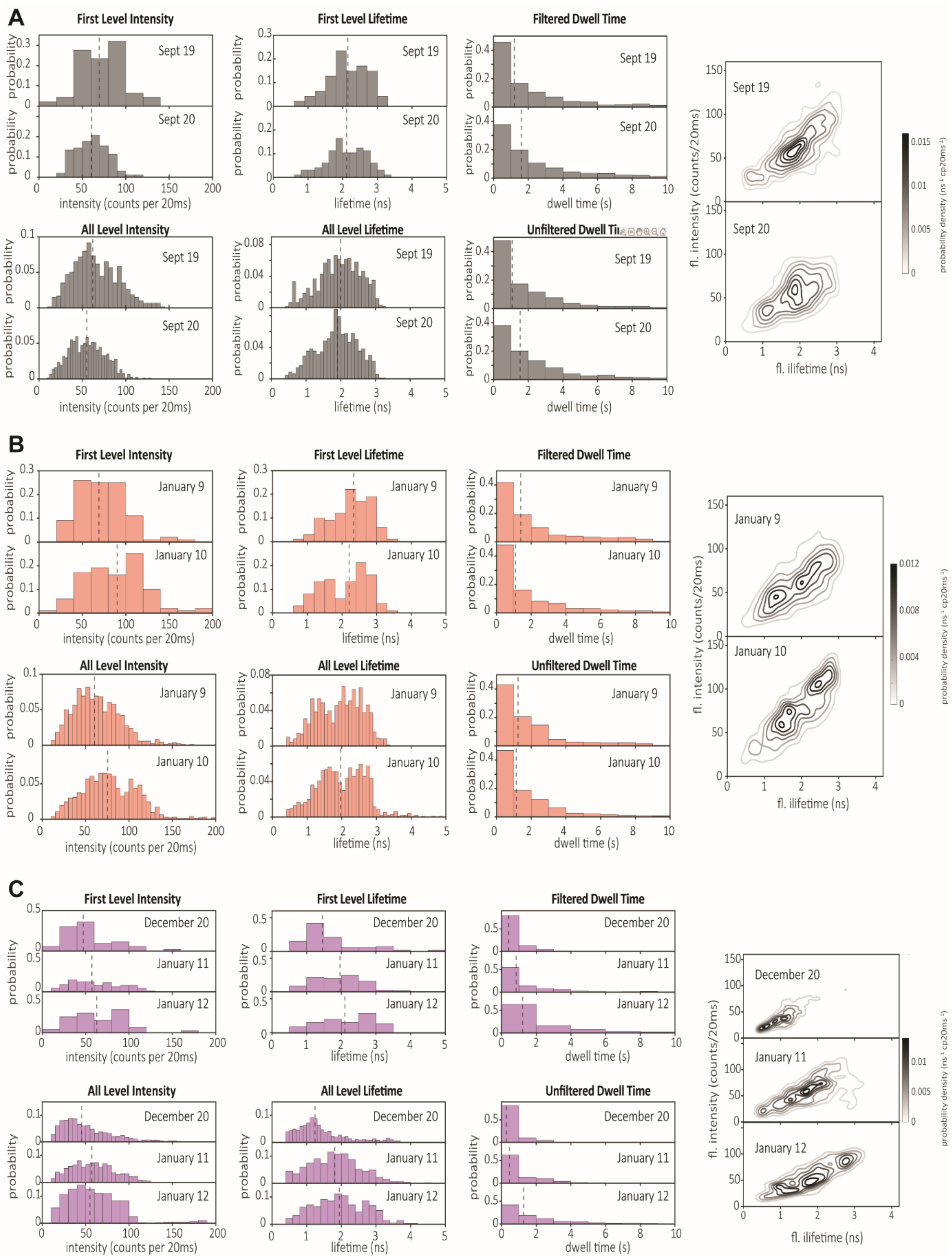


**Figure S14. Day-to-day variability.**

Histograms from each day of single-molecule data collections of LHCII from WT (A), LHCII-qH(−) (B) and LHCII-qH(+) (C) for first and all level intensity, lifetime, filtered and unfiltered dwell times and density plots of intensity and lifetime from filtered states. Medians shown as dashed lines in histogram.

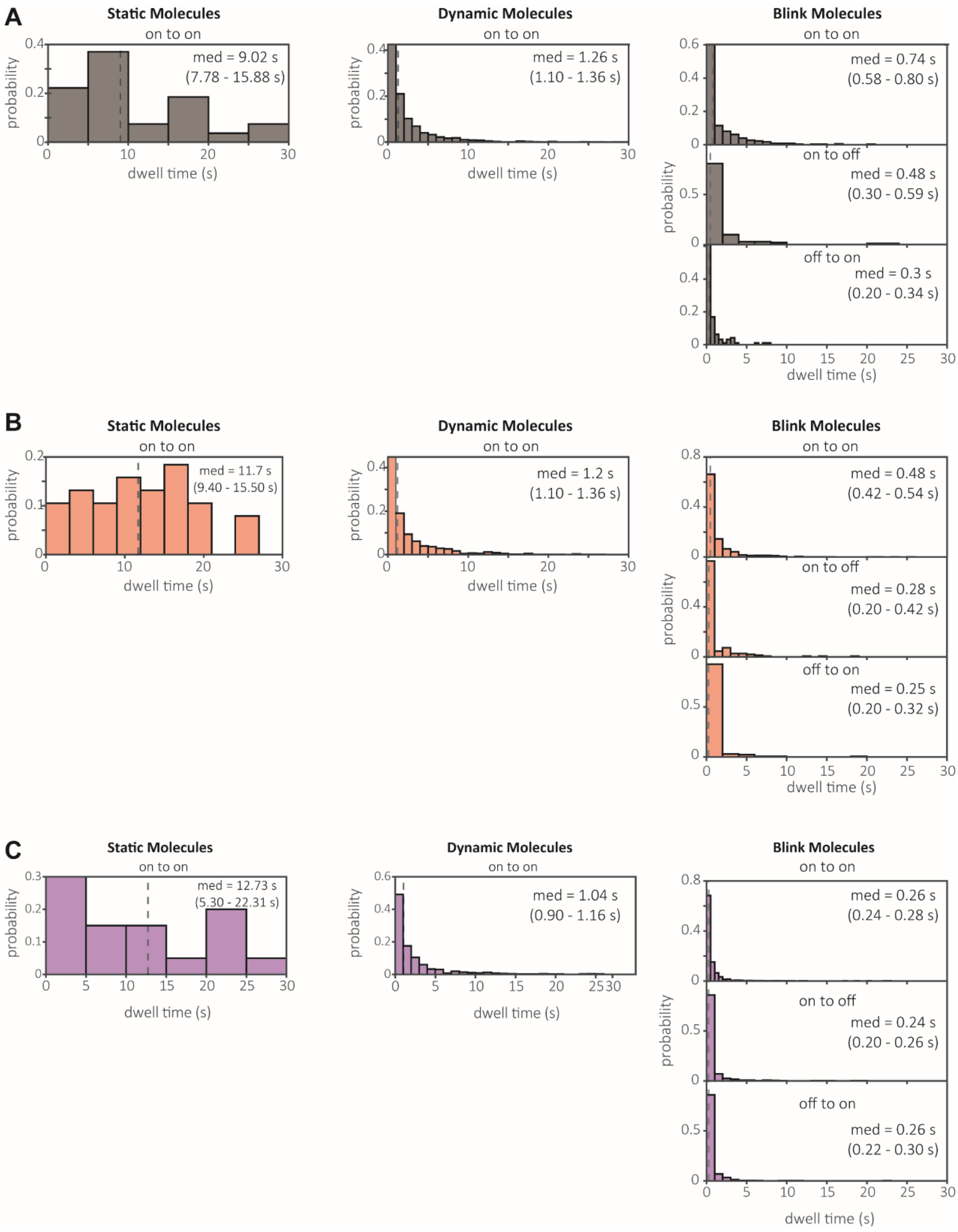


**Figure S15. Switching kinetics by transition type.**

Unfiltered dwell times by transition type for LHCII from WT (A), LHCII-qH(−) (B) and LHCII-qH(+) (C) molecules. On-to-on is the dwell time of a molecule in an on state before switching to another on state. On-to-off is the dwell time of a molecule in an on state before switching to an off state. Off-to-on is the dwell time of a molecule in an off state before switching to an on state. Medians shown by dashed line. Median values of on-to-on in static and dynamic molecules differ slightly from those in the main text because only states before a transition were considered, therefore the last state of every molecule is not included in these histograms. 95% confidence intervals were determined by the median of 1000 bootstrapped samples and are shown between brackets.

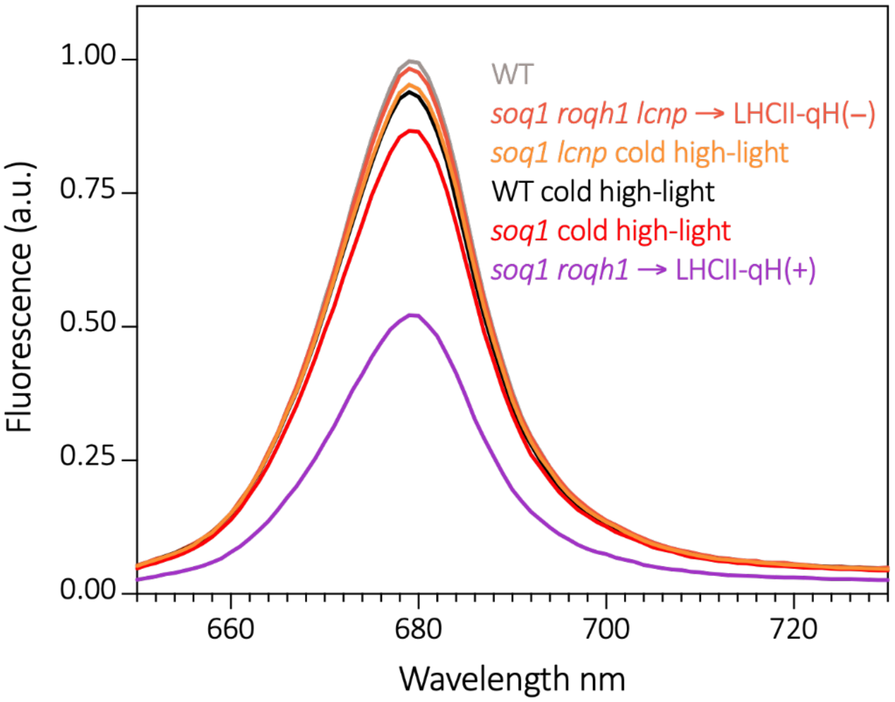


**Figure S16. Room temperature fluorescence of LHCII trimers purified in β-DDM detergent.**

LHCII trimers were isolated in n-dodecyl β-D-maltoside (β-DDM). The fluorescence yield is similar to the samples purified in α-DDM (see Figure 1), with LHCII-qH(+) from *soq1 roqh1* displaying a yield around 53 ± 1% of controls. Excitation was at 625 nm, and data normalized to the maximum OD in the Qy peak and maximum fluorescence of WT LHCII. Data represent mean of three replicates of different LHCII purifications from the same thylakoid batches (two for WT).
